# Deep learning of fossil pollen morphology reveals 25,000 years of ecological change in East African grasslands

**DOI:** 10.1101/2024.09.23.612957

**Authors:** Marc-Élie Adaimé, Shu Kong, Michael A. Urban, F. Alayne Street-Perrott, Dirk Verschuren, Surangi W. Punyasena

## Abstract

Grass pollen is largely overlooked in investigating grassland evolution because the pollen of most species cannot be differentiated using traditional optical microscopy. However, deep learning can quantify small variations in pollen morphology visible under superresolution microscopy. We use the abstract features output by deep learning to estimate the taxonomic diversity and physiology of fossil grass pollen assemblages. Using a semi-supervised learning strategy, we trained convolutional neural networks (CNNs) on superresolution pollen images of modern grasses and unlabeled fossil Poaceae. Our models captured features that reflected both the taxonomic diversity of grass communities along an elevational gradient and morphological differences between C_3_ and C_4_ species. We applied our trained models to fossil grass pollen assemblages from a 25,000-year lake-sediment record from eastern equatorial Africa (Mt. Kenya) and correlated past shifts in grass diversity with atmospheric CO_2_ concentration and proxy records of local temperature, precipitation, and fire occurrence. We quantified changes in grass diversity using morphological variability of fossil pollen assemblages, approximated by the Shannon entropy of CNN features. Our data show that grassland species diversity was strongly reduced between 21,500 and 16,000 years ago, coincident with most severe regional cooling during the last ice age. C_3_:C_4_ ratios reconstructed using a gradient-boosted decision tree classifier infer a gradual decrease in C_4_ grasses since the late-glacial to Holocene transition, associated with decreasing fire activity and elevated temperatures. Our results demonstrate that CNN features of pollen morphology can advance palynological analysis, enabling robust estimation of grass diversity and C_3_:C_4_ ratio in ancient grassland ecosystems.

**Significance:** Although the pollen of most grass species are morphologically indistinguishable using traditional optical microscopy, we show that they can be differentiated through deep learning analyses of superresolution images. Abstracted morphological features derived from convolutional neural networks can be used to quantify the biological and physiological diversity of grass pollen assemblages, without *a priori* knowledge of the species present, and used to reconstruct past changes in the taxonomic diversity and relative abundance of C_4_ grasses in ancient grasslands. This approach unlocks ecological information previously unattainable from the fossil pollen record and demonstrates that deep learning can solve some of the most intractable identification problems in the reconstruction of past vegetation dynamics.

## Introduction

The evolutionary and ecological history of grasses and grasslands is recorded in the global fossil pollen record. However, despite its abundance, grass pollen is underutilized in paleoecological analyses because the pollen of Poaceae species tends to be visually indistinguishable using traditional optical microscopy. This vast paleontological record has been largely limited to estimating past changes in grass abundance (1, 2). As a result, the fossil record of grass pollen has contributed little insight into the evolution of the ∼11,800 species of this diverse plant lineage (3) or the establishment and community dynamics of grassland ecosystems.

Grasslands are a relatively recent ecosystem in Earth history, first appearing in the Eocene-Oligocene (4). Today they cover ∼40% of Earth’s terrestrial surface and contribute substantially to ecosystem services such as soil fertility, carbon storage, and biosphere control of atmospheric greenhouse gas concentrations (5, 6). Grasslands are characterized by unique ecological properties, including high water-use efficiency and high albedo, and play a critical role in regulating the global biogeochemical cycles of nitrogen, silica, and carbon (5, 7). The productivity and ecological resilience of grasslands is positively correlated with their diversity (8), with high species richness promoting resilience to environmental stressors (8, 9).

Grass species employ one of two photosynthetic pathways, C_3_ and C_4_. C_4_ photosynthesis uses water more efficiently and is more tolerant of low atmospheric CO_2_ than the ancestral C_3_ pathway (10). C_4_ photosynthesis evolved independently in multiple grass lineages, allowing these species to eventually dominate the world’s grasslands by the late Miocene (ca. 8 million years ago) (4, 11, 12). The global expansion of C_4_ grasslands during the late Cenozoic occurred alongside a downward trend in atmospheric CO_2_ concentration, associated with major environmental changes such as the growth of continental ice sheets and the increased prevalence of seasonally dry tropical and temperate climate regimes (4, 13). During the Quaternary period (the last 2.5 million years), grassland ecosystems expanded throughout the tropics due to the generally cool and dry climatic conditions characterizing Pleistocene glacial periods in low-latitude regions (14). Pollen records from southern and southeastern Brazil indicate that now forested regions were dominated by grasslands during glacial times (15). Similarly, in Africa, grasslands expanded during the Last Glacial Maximum (LGM) (16).

Sensitivity to climate, combined with their widespread geographic distribution and extensive fossil record, make grasses an important paleobiological proxy in the reconstruction of past climates and environments (17, 18). However, interpreting the fossil record of grasses has been a long-standing challenge. Fossil Poaceae leaves and stems are rarely preserved with enough morphological detail to distinguish them from other monocotyledonous plants (4). Fossil phytoliths have generally provided the most information about the evolution of open grasslands in contemporary paleoecological research (4, 12). However, multiple phytolith morphotypes are shared between C_3_ and C_4_ species, so phytoliths alone cannot directly capture taxonomic and physiological diversity (12, 19). Multiproxy approaches combining pollen, starch grains, and phytoliths have advanced identification of fossil grasses in archeobotanical contexts (20), and Fourier-transform infrared (FTIR) microspectroscopy can classify modern Poaceae pollen based on their chemical spectra (21). However, starch grain analysis and FTIR are not generally applicable to community-level or deep-time (≥2 Ma) paleoecology. Furthermore, starch and phytolith evidence are context- and tissue-dependent, while FTIR classifications require specific sample preparation, rely on sporopollenin chemical spectra that can be altered by sedimentation processes, and have thus far been applied to only a limited number of taxa (21).

Fossil grass pollen has been largely ignored as a source of paleoecological information because grass pollen grains are morphologically similar across the family. All have a single annulated pore and a psilate (visually smooth) outer exine surface. However, small variations in the patterning of the pollen grain surface visible under electron microscopy have been shown to be taxonomically meaningful and can be characterized quantitatively (22, 23, 2, 24–26). While pollen size alone is insufficient to tell apart grasses, there are also quantifiable differences in size among species (27). However, all of these previous categorizations of grass pollen morphotypes have been too broad to capture changes in the taxonomic and physiological composition of grassland communities on millennial or longer timescales.

New approaches are therefore needed to effectively reconstruct the diversity and composition of fossil grasslands. Here we propose that morphological differences among the pollen of different grass species revealed in superresolution microscopic images, though subtle, can provide the basis for training deep learning models. Machine learning has become an important component of the palynological toolkit over the last decade (e.g., 28–30), but with few exceptions (31–34) it has been used to automate, rather than to improve upon, pollen identifications. In this study, we developed deep learning pipelines that use convolutional neural networks (CNNs) trained on superresolution images (0.04 µm resolution) of modern and fossil Poaceae pollen (Fig. 1A), with the goal of improving on expert classifications of grass pollen. The abstract features derived from these CNNs quantify morphological differences that capture the taxonomic and physiological diversity of grass pollen assemblages. Our first computational pipeline quantifies the taxonomic diversity in fossil pollen assemblages by using Shannon entropy to approximate morphological variability (Fig. 1B). Our second pipeline uses a gradient-boosted decision tree classifier (35, 36) to identify the variation within the CNN-derived features of modern Poaceae pollen that are most strongly associated with either C_3_ or C_4_ grass species.

**Fig. 1.**
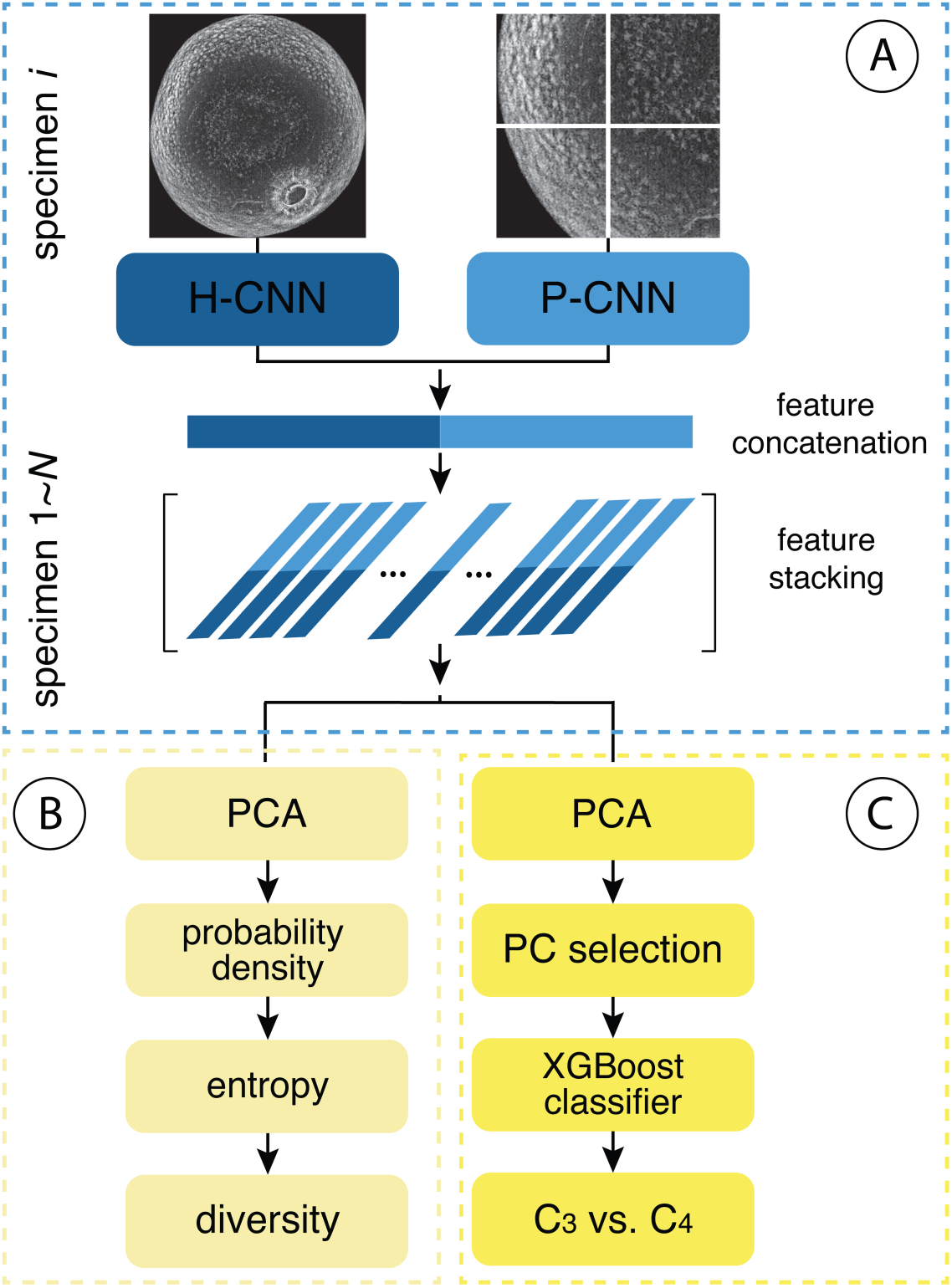
Flowchart illustrating the pipeline of computational analyses performed in this study. (A) Two superresolution microscopic representations of a grass pollen grain (the maximum intensity projection of the image stack and overlapping crops of the maximum intensity projection) are passed to convolutional neural networks (CNNs) trained to recognize the holistic morphology (H-CNN) and surface texture of patches (P-CNN) of grass pollen. The morphological features extracted by both CNNs are concatenated to form a comprehensive feature vector representing each specimen. These vectors are then stacked across all *N* grass pollen grains present in an assemblage, forming a feature matrix of all the concatenated feature vectors. (B) In our assessment of grass diversity, the PCA scores of the feature matrix are used to estimate probability densities for pollen assemblages using kernel density estimation. Shannon entropy is calculated over these estimated densities to provide a quantitative measure of diversity. (C) In our assessment of C_3_:C_4_ ratios, we apply PCA to the feature matrix and pass the most informative principal components to a gradient boosted decision tree (XGBoost) trained to discriminate between C_3_ and C_4_ grasses.

We then applied our trained CNN models to superresolution images of fossil grass pollen assemblages extracted from 24 time windows in the 25,000-year sediment record of Lake Rutundu (0.0416° S, 37.4638° E), a crater lake located at 3078 m elevation in the heathland-moorland (‘subalpine’) zone of Mt. Kenya in eastern equatorial Africa (37) (Fig. 2). We reconstructed the taxonomic diversity and C_4_ fraction of the high-mountain grasslands surrounding Lake Rutundu relative to changes in local environmental factors such as fire regime (38), temperature (39), and precipitation (40), as well as the known evolution of atmospheric CO_2_ concentration (41). The wealth of paleoecological and paleoclimate data available from previous research on the Lake Rutundu record (37, 38, 42, 43) allowed direct comparison of our results with earlier reconstructions of the history of grasslands on Mt. Kenya. Our findings demonstrate that the combination of superresolution imaging and deep learning enables – for the first time – the estimation of past grass diversity and C_3_:C_4_ ratios from fossil pollen morphology. With these tools, we can interpret the diversity and composition of ancient grass communities even in the absence of species identifications.

**Fig. 2.**
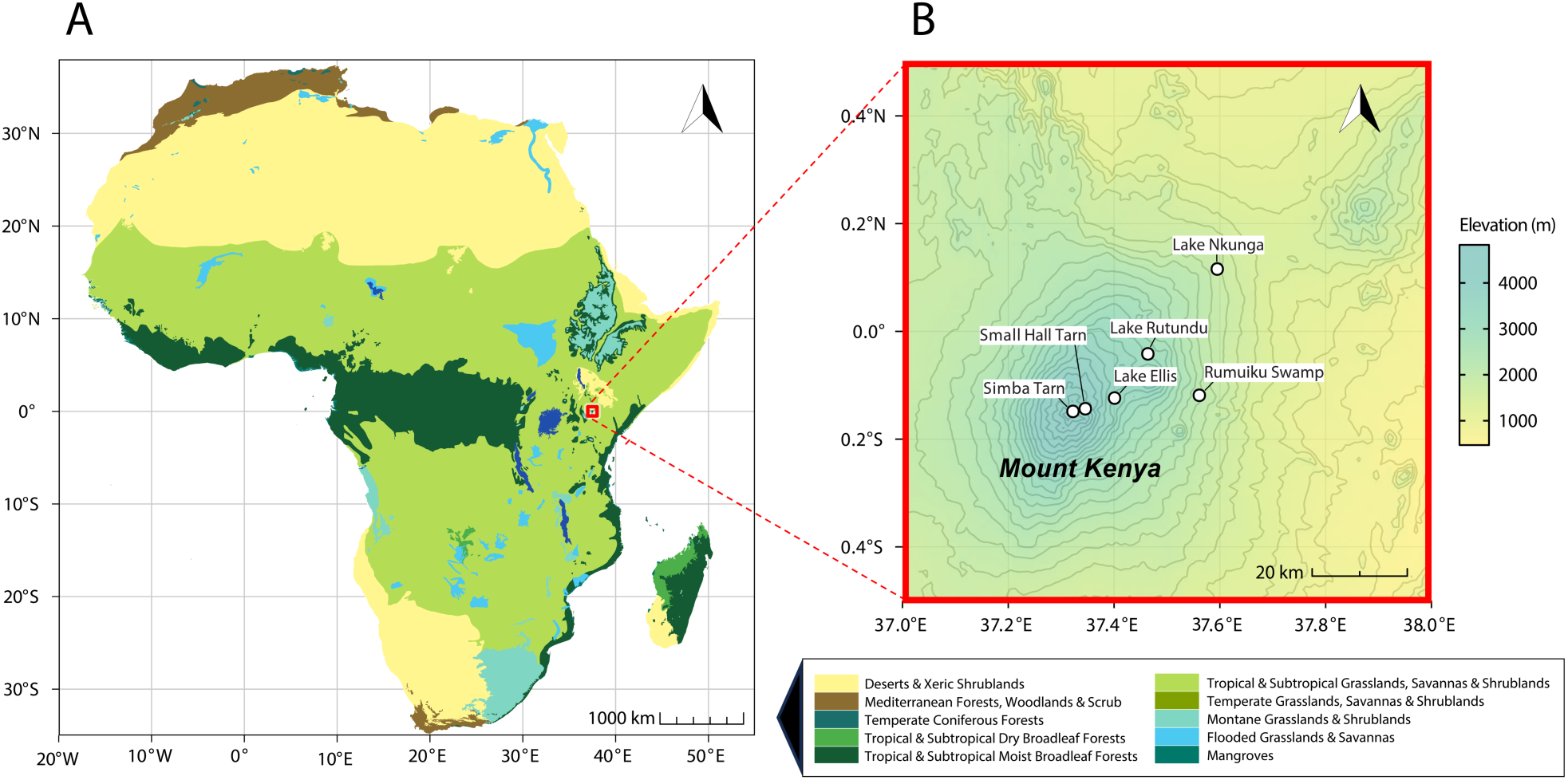
Setting of the 25,000-year grass-pollen record used for method validation. (A) Distribution of major vegetation biomes across the African continent, with the location of Mt. Kenya in eastern equatorial Africa highlighted by a red square. (B) Elevation gradient in the Mt. Kenya region with the locations of our six sources of fossil grass pollen: the 25,000-year sediment record from Lake Rutundu, and the five additional lakes (Lake Nkunga, Rumuiku Swamp, Lake Ellis, Small Hall Tarn, and Simba Tarn) located between 1820 m and 4585 m elevation where we extracted subfossil pollen from recently deposited sediment.

## Results

### CNNs can discriminate the pollen of different grass species

We trained two separate K-way classification CNNs in a semi-supervised learning framework, using superresolution structured illumination (SR-SIM) (44) images of 1,241 modern pollen grains identified to species and 1,503 unlabeled fossil pollen grains. Modern pollen grains were isolated from herbarium specimens of 60 extant Poaceae species (3–37 specimens per species; SI Appendix, Tables S1 and S2). Fossil material was isolated from the principal Lake Rutundu sediment core (37) and from recently deposited sediments in five other lakes on Mt. Kenya (SI Appendix, Tables S3 and S4). The first CNN (holistic CNN; H-CNN) was trained on maximum intensity projections (MIPs) of whole grains. The second CNN (patch CNN; P-CNN) used small, cropped segments of the MIP for training (Fig. 1A). CNNs are a series of convolutional and pooling layers with a final classification layer that produces probabilities of class labels (45). Notably, we employed CNNs to extract abstracted morphological characters for downstream analyses and not for classification. Our analysis ignores the final classification layer and instead uses the penultimate layer containing the highest-level representation of a specimen’s morphology, or “features” (33). However, we assessed the quality of these CNN features by calculating the average classification accuracy of our trained models (SI Appendix, Fig. S1). The H-CNN and P-CNN achieved an average accuracy of 61.40% and 72.51%, respectively (SI Appendix, Table S5). Fusion of the two CNNs achieved an average accuracy of 73.95% (SI Appendix, Table S5). All but three species had ≥19 specimens for training. *Andropogon chrysostachyus*, with only 3 specimens for training, was never identified correctly (i.e., 0% accuracy). All other species had average accuracies >23%, which is significantly greater than chance (1.67%).

### Morphological variation within and among grass pollen is taxonomically meaningful and reflects species diversity

The CNN-derived morphological features of grass pollen show clear taxonomic structure. Species belonging to the same genus cluster significantly closer together in CNN feature space than species from different genera (p = 0.0001), indicating that the learned features capture biologically meaningful variation rather than overfitting to individual species (SI Appendix, Fig. S2A). We also found that morphological variability within species is not significantly affected by whether the grains came from a single or from multiple individuals. The mean within-species variance did not differ between these two groups (p = 0.498), suggesting that intra-individual and inter-individual morphological variability are broadly similar (SI Appendix, Fig. S2B).

We next used artificial pollen assemblages (i.e., simulated grass communities) to test whether the Shannon entropy (46) of the CNN features of a fossil grass pollen assemblage captured its taxonomic diversity. We randomly split our modern Poaceae dataset into 30 species used to train the CNNs and 30 “unseen” species. We randomly drew 50 grains from the 30 “unseen” species 100 times to form 100 artificial pollen assemblages of varying species richness and relative abundance. We forward-passed these grains through our trained CNN models and calculated the assemblages’ Shannon entropy from the distribution of their CNN features (see Methods). We used PCA to reduce the dimensionality of our CNN feature matrix, producing three PC subspaces of Shannon entropy identified as PC1, PC1-PC2, and PC1-PC2-PC3. There was a strong positive correlation between the Shannon entropy of the CNN features and the Shannon diversity index (46) of the artificial pollen assemblages (r = 0.47, 0.58, 0.66 for each respective PCA subspace; p < 0.0001) (SI Appendix, Fig. S3). The Shannon index is a standard statistical measure of diversity that accounts for both species richness and evenness (47). These results demonstrate that the Shannon entropy of CNN features effectively captures variation in species diversity.

### Modern lake sediment samples capture modern grass diversity gradients

We used grass pollen isolated from recently deposited sediments of six Mt. Kenya lakes situated between 1820 and 4585 m elevation (equivalent to a 16 °C difference in mean annual temperature [MAT]; ref. 48) to test whether the Shannon entropy of CNN features captures modern elevational gradients in grass diversity. Using the first principal component (PC1) of our CNN feature matrix as the diversity metric, we evaluated variation in diversity across a set of kernel bandwidth multipliers (Fig. S4). Leave-one-out cross-validation (LOO-CV) (49) identified h^∗^ ≈ 3.87, ℎ^∗^ ≈ 5.99, and ℎ^∗^ ≈ 6.24 as the optimal bandwidths for the PC1, PC1-PC2, and PC1-PC2-PC3 subspaces. At smoothing (ℎ) values below the optimal bandwidth, the PC1 subspace recovered a mid-elevational (∼3000–3500 m a.s.l.) peak in diversity with lower diversity both at lower and higher elevations (SI Appendix, Fig. S4). This closely matches data from vegetation surveys, which indicate that maximum grass species richness on Mt. Kenya (9–11 species) occurs in the heathland-moorland zone between 3100 and 3400 m altitude (where present-day MAT is *ca.* 8 °C; ref. 38), with lower richness both in low-elevation savanna woodland (1–4 species) and in the upper alpine zone above 4000 m (1–5 species) (50). With greater smoothing (where ℎ = αℎ^∗^, α = √2), between-site differences in PC1 are reduced, resulting in weak expression of the mid-elevation diversity maximum (Fig. S4).

### Changes in grass diversity on Mt. Kenya through time

We applied the same approach (using the PCA scores of the Shannon entropy of the CNN features of a pollen assemblage as a proxy for grass diversity) to fossil grass pollen assemblages isolated from 24 more or less equally spaced time windows in the 25,000-year Lake Rutundu sediment record. We calculated entropy for each PCA subspace at their optimal bandwidths (ℎ^∗^), as derived from the modern-day elevation gradient in grass species diversity (Fig. S4), and compared the long-term temporal trends in entropy (i.e., the inferred past grass diversity) with time series of environmental variables potentially forcing these trends (Fig. 3C-D).

**Fig. 3.**
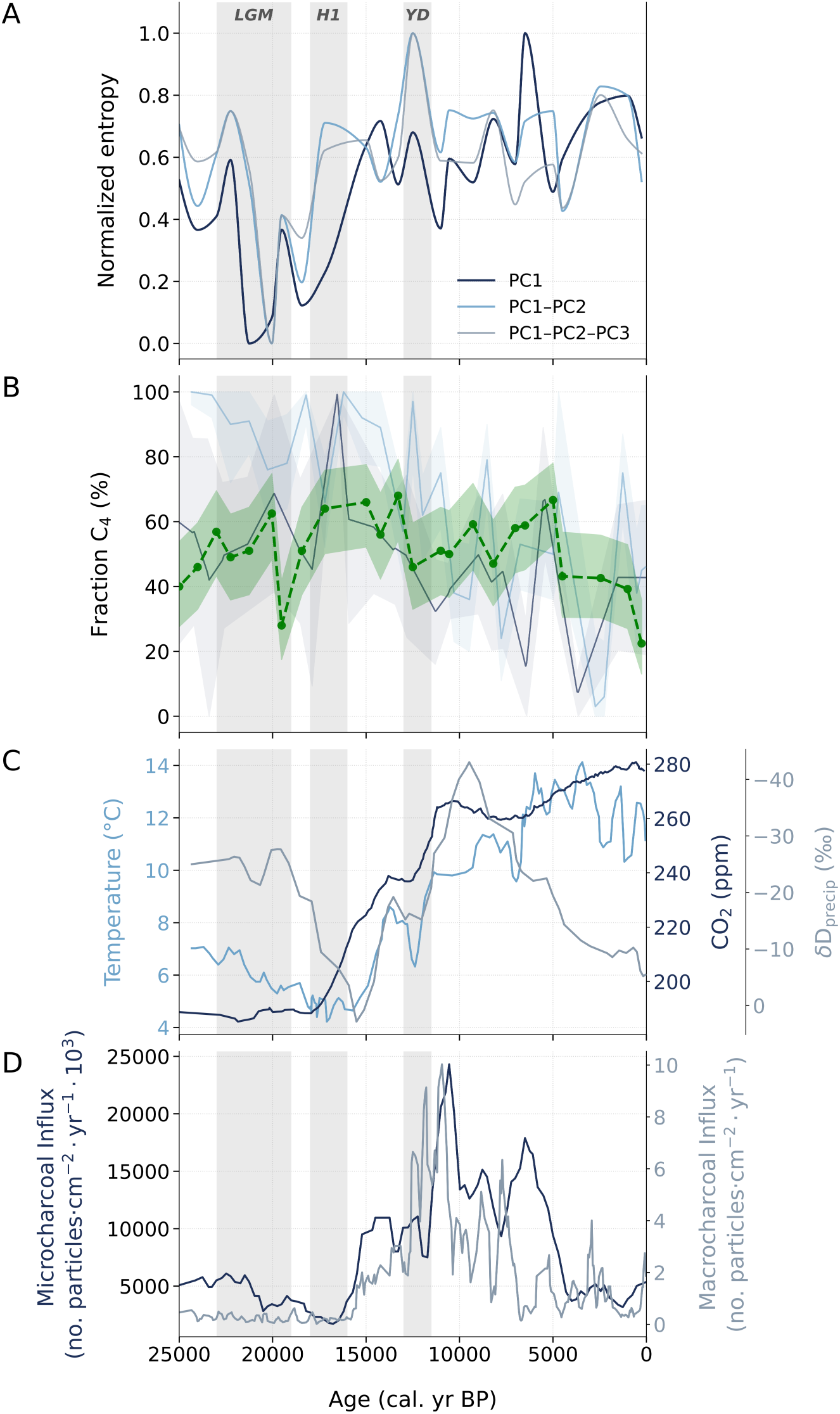
Changes in grass diversity and proportion of C_4_ grasses in subalpine grasslands of Mt. Kenya over the past 25,000 years in relation to paleoenvironmental history. Vertical gray shading delineates major late-glacial climatic intervals: the Last Glacial Maximum (LGM; 23,000–19,000 cal. yr BP), Heinrich Stadial 1 (H1; 18,000–16,000 cal. yr BP), and the Younger Dryas (YD; 13,000–11,500 cal. yr BP). Paleoenvironmental proxies (C, D) were interpolated using moving averages with a window size of 4. (A) Grass species richness as inferred from the morphological diversity of grass pollen, as estimated by differential entropy of Gaussian kernel density estimates (KDEs) on PCA scores (PC1, PC1-PC2, and PC1-PC2-PC3). Bandwidth selection, curve smoothing, and evaluation are detailed in Methods. Results are normalized for visualization. (B) Proportion of C_4_ grasses of the grassland surrounding Lake Rutundu as estimated by this study (dashed green line), the carbon-isotopic signature (δ^13^C) of individual grass pollen grains (solid blue line; ref. 38), and fossil grass cuticle (solid purple line; ref. 37). Uncertainty is shown as shaded bands: 95% Wilson confidence intervals for this study, 95% credible intervals from the Bayesian mixture model in (38), and bounds from “certain only” versus “certain + likely” C_4_ cuticle identifications in (37). (C) Antarctic ice-core record of atmospheric CO_2_ concentration (41) and reconstructions of local mean annual temperature (MAT) and precipitation derived from, respectively, bacterial membrane lipids and the hydrogen-isotopic signature (δD_precip_) of plant leaf-wax alkanes extracted from Lake Rutundu sediments (28, 29). More depleted δD_precip_ values indicate wetter conditions. (D) History of fire prevalence on Mt. Kenya reconstructed from estimates of biomass burning based on sedimentary fluxes of micro- and macrocharcoal (38).

We find that grass diversity in the subalpine grasslands on Mt. Kenya was strongly reduced during the LGM and earliest late-glacial period (*ca.* 21,500–16,000 years ago), broadly coeval with the period of most strongly reduced MAT in eastern equatorial Africa (51) and at high elevations on Mt. Kenya in particular (7–8 °C cooler than today; ref. 38) (Fig. 3A, C). Grass diversity has been substantially higher from *ca.* 15,000 years ago until the present, and was also high under the climate regime immediately before the LGM when glacial cooling was less extreme (4–6 °C cooler than today; ref. 38). Entropy values for all three PCA subspaces fluctuate throughout the Holocene period but with no sustained trend (Fig. 3A), suggesting no or limited change in the diversity of Mt. Kenya grasslands at the elevation of Lake Rutundu during this time period. Over the 25,000-year record as a whole, entropy values for subspaces PC1 and PC1-PC2 consistently and positively correlated with both changes in atmospheric CO_2_ and local temperature, although the strength of the relationships varied. For PC1, r = 0.67 for CO_2_ and 0.68 for temperature (p < 0.001); for PC1-PC2, r = 0.47 for CO_2_ and 0.48 for temperature (p < 0.05) (Fig. 3A and SI Appendix, Fig. S5). Entropy also correlated significantly with fire regime proxies (micro- and macrocharcoal influx) in the morphological subspace represented by PC1-PC2 (r = 0.45 for microcharcoal and 0.44 for macrocharcoal, p < 0.05), while they were weak and not significant for PC1. No diversity estimates correlated with reconstructed precipitation and no significant relationships were detected in the PC1-PC2-PC3 subspace (Fig. 3A and SI Appendix, Fig. S5).

The CNN features of each Rutundu fossil grass pollen assemblage also capture the morphological disparity among the assemblages’ grass pollen grains. This disparity reflects the underlying species composition. Therefore, changes in the morphological space occupied by the CNN features likely represent changes in the taxonomic composition of grass communities on Mt. Kenya through time. We used PCA to reduce the dimensionality of the dataset and to visualize the distribution of CNN features in each time window. We found that the combined morphological diversity of all 24 fossil samples is contained within the diversity present in our reference dataset of 60 modern grass species (Fig. 4), suggesting that our reference data capture the majority of the morphological disparity within our fossil dataset. However, the mean distribution of CNN features in the fossil dataset is significantly different from that in the modern dataset for PC1 (Welch’s t-test, p < 0.0001). Moreover, the morphological space occupied by the fossil assemblages varies through time. For example, the four assemblages dated to the LGM (21,260–18,435 cal yr. BP) are distinct from those of other time windows in that they occupy a distinctly smaller area in the morphospace defined by PC1 and PC2 (Fig. 4).

**Fig. 4.**
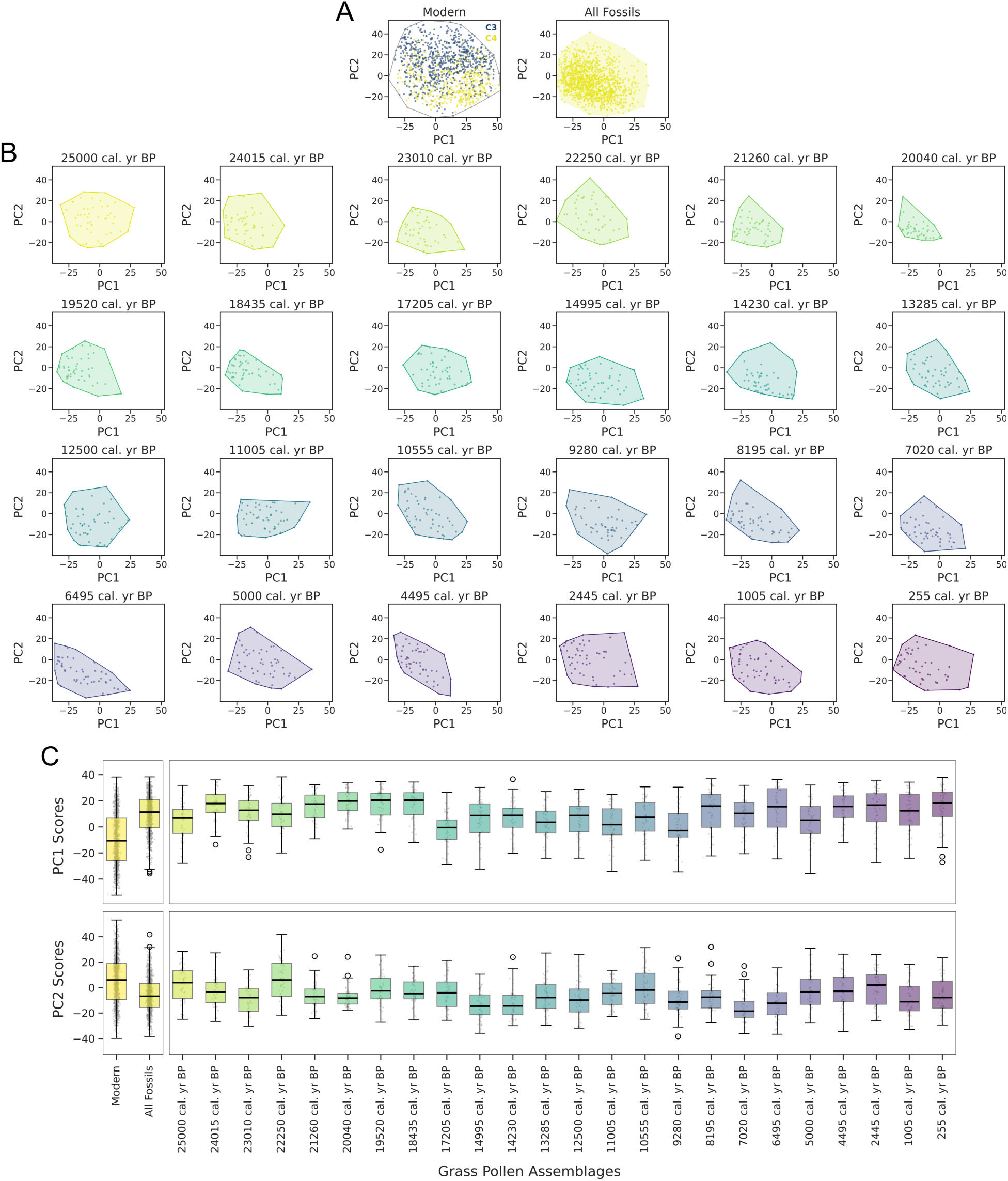
(A) PC1-PC2 biplots of the morphological variation among 1,241 grass pollen grains in our 60-species modern dataset and the projected placement of 1,204 fossil grains from all 24 analyzed time windows in the Lake Rutundu record. There is partial separation between the pollen of modern C_3_ (blue) and C_4_ (yellow) grass species, and the morphological variation among the fossil Poaceae pollen falls largely within a subset of the modern range of morphospace. PC1 and PC2 together account for 17.05% of the variance. (B) PC1-PC2 biplots of the pollen morphological variation in fossil assemblages from each analyzed time window, with shaded convex hulls delineating the morphological range present in each assemblage. The colors for each time window match the boxplots below. (C) Boxplots of the median values and ranges of PC1 and PC2 scores of the 24 fossil assemblages, highlighting the distinctively low PC1 scores of the four assemblages dated to between 21,160 and 18,435 BP.

### Grass pollen morphology captures photosynthetic pathway

To determine whether there are consistent morphological differences between the pollen of C_3_ and C_4_ grass species, we averaged the CNN features of our modern grass pollen dataset by species and used PCA to reduce its dimensionality. The first two principal components significantly discriminated between the pollen of C_3_ and C_4_ species (Welch’s t-test, p = 0.0075 for PC1 and p < 0.001 for PC2) (Figs. 4, 5D). However, tests for phylogenetic signal revealed strong evolutionary structure in both PC1 (λ = 0.854, p < 0.001) and PC2 (λ = 0.738, p < 0.001), indicating that morphological traits may also be strongly influenced by evolutionary relatedness. When corrected for phylogenetic relatedness using Phylogenetic Generalized Least Squares (PGLS) regression (see Methods), the difference in mean PC1 values between C_3_ and C_4_ species remains significant (p = 0.014, λ = 0.906), but the difference in mean PC2 values does not (p = 0.470, λ = 0.706).

**Fig. 5.**
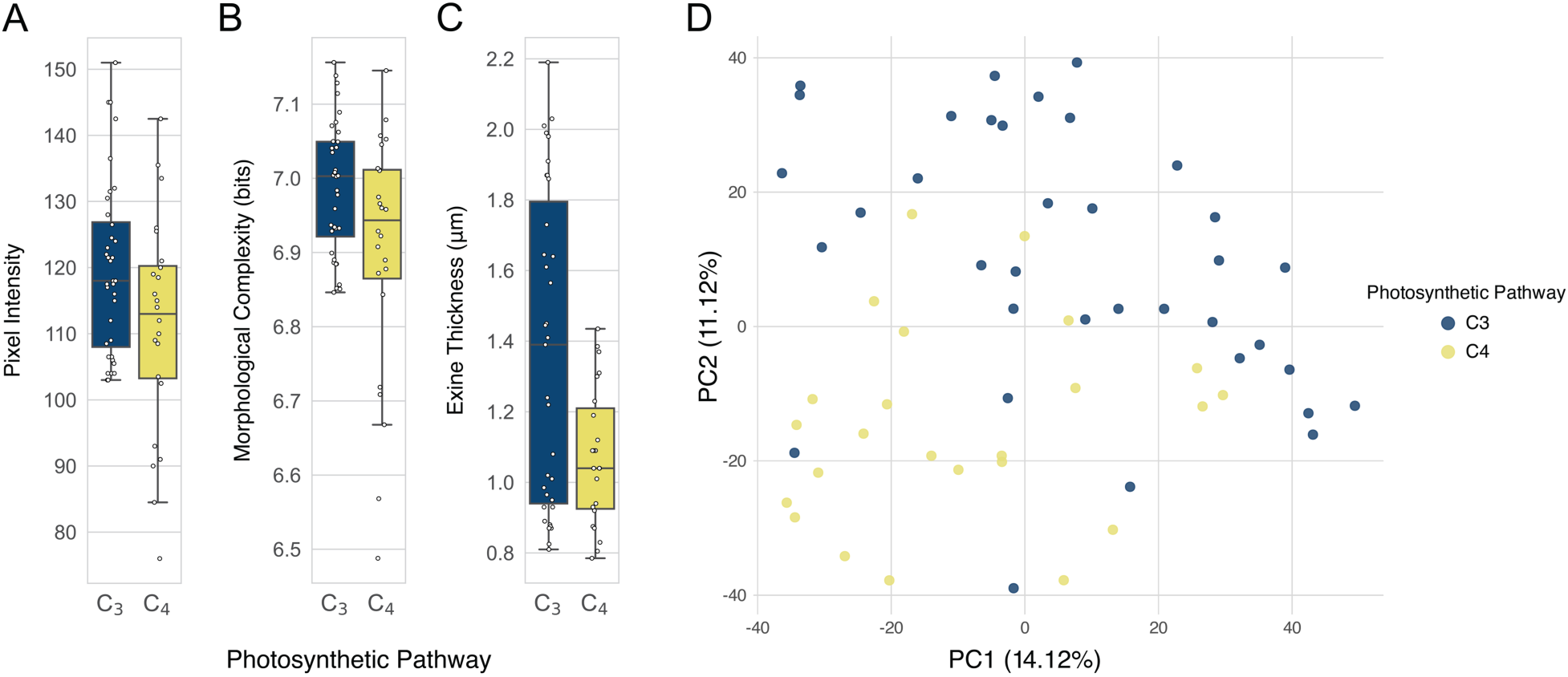
Measurements by this study and previous morphometric analyses showing that C_3_ grass pollen has a thicker exine wall and more complex surface patterning than C_4_ grass pollen. Boxplots showing the difference in (A) pixel intensity (brightness) and (B) morphological complexity (measured using Shannon entropy) between pollen from 36 C_3_ and 24 C_4_ grass species in our modern reference dataset, and (C) in exine thickness (µm) of pollen from 35 C_3_ and 23 C_4_ grass species determined by (24). (D) PC1-PC2 biplot of mean CNN-derived pollen morphological features across 60 modern Poaceae species illustrating the morphological differentiation between C_3_ and C_4_ grasses. Each point represents a mean species-level feature vector.

While we cannot link variation in PC1 to specific pollen morphological differences between C_3_ and C_4_ grasses, the median pixel brightness of specimen images was found to be higher in C_3_ grasses (Fig. 5A; Welch’s t-test, p = 0.029), which suggests that they have a thicker exine wall, on average, than the pollen of C_4_ grasses. The entropy of the distribution of pixel values was also higher in C_3_ grass pollen (Fig. 5B; Welch’s t-test, p = 0.041), suggesting a more complex surface texture (note that this measure is distinct from our calculation of Shannon entropy used for diversity estimation). Using actual measurements of exine thickness in an independent dataset of 35 C_3_ and 23 C_4_ grass species assembled by (24), we tested the hypothesis that pollen grains of C_3_ grasses have thicker exine walls and found this to be the case (Fig. 5C; Welch’s t-test, p = 0.001).

### Late Quaternary changes in C_4_ grass fraction on Mt. Kenya

Given the morphological differences between pollen of living C_3_ and C_4_ grasses, we finally attempted to reconstruct changes through time in the fraction of C_4_ grasses on Mt. Kenya as registered in the Lake Rutundu record. We first trained a C_3_ versus C_4_ classifier on the CNN features of all 1,241 individual modern grass pollen grains. We used PCA to reduce the dimensionality of the feature matrix and applied a gradient-boosted decision tree classifier to the first 230 principal components, which together represent 95% of the explained variance in pollen morphology. The classifier achieved high prediction accuracy in distinguishing C_3_ and C_4_ grains, confirmed by cross-validation (µ = 91.1%, σ = 2.5%).

We next projected the CNN features of the 1,204 fossil pollen grains from Lake Rutundu onto the PCA space defined by the CNN features of the modern grass pollen (see Methods). We forward-passed the first 230 principal components through the trained gradient-boosted decision tree classifier to determine the proportion of pollen derived from C_4_ grasses in each of the 24 fossil assemblages of the Rutundu record. The C_4_ fraction ranged between 22–68%, with a general increase from ∼40% *ca.* 25,000 years ago to 50–60% during the LGM (with the exception of one outlying value of 28% *ca.* 19,500 years ago), followed by an overall decreasing trend from the late-glacial period *ca*. 15,000–13,000 years ago until the late-Holocene and present (Fig. 3B). These results are consistent with estimates of C_4_ grass fraction inferred from taxonomically diagnostic grass cuticle fragments preserved in the same Lake Rutundu record, which range between 16–73% and show a stable fraction of 50–60% C_4_ grasses prior to and during the LGM, followed by a general decline over the past 15,000 years, albeit punctuated by large fluctuations (Fig. 3B) (37). They are less in agreement with a reconstruction of C_4_ grass abundance based on the carbon isotope ratio (δ¹³C) of fossil grass pollen from Lake Rutundu, which also shows a long-term decline in the proportion of C_4_ grasses during the postglacial period, but infers values of 75–100% before and during the LGM (Fig. 3B) (38). Nevertheless, pairwise correlations between these three C_4_ data series, interpolated using varying degrees of spline smoothing, are all highly significant (p < 0.0001) (SI Appendix, Fig. S6), indicating that they all infer a broadly similar long-term trend, namely the general decline of C_4_ grasses relative to C_3_ grasses during the postglacial period on Mt. Kenya. This reconstructed temporal trend in C_4_ grass abundance on Mt. Kenya is negatively related to past changes in atmospheric CO_2_ and temperature (Fig. 3C) and positively related to fire prevalence (Fig. 3D), although neither of these correlations is significant (p > 0.05 in all cases).

## Discussion

Deep learning techniques have allowed us to quantify the sub-micrometer scale differences in pollen morphology made visible by superresolution microscopy, and to statistically differentiate different morphotypes of grass (Poaceae) pollen. With these computational tools, we demonstrate the capability to more consistently discriminate among the pollen of known grass species (SI Appendix, Fig. S1), infer the diversity (species richness) of grass communities from pollen morphological diversity (Fig. 3A and SI Appendix, Figs. S3–S5), and capture past changes in the relative abundance of C_3_ and C_4_ grasses (Fig. 3B and SI Appendix, Fig. S5). In addition to opening important new opportunities in paleobotanical research, our results also support the important assumption in palynology that the morphological disparity among pollen grains from a single plant specimen is smaller than that between species (52) and provides a meaningful representation of intraspecific pollen variability. Indeed, the variation in CNN features of pollen from multiple plants within a single species was found to not differ significantly from that of a single plant, and pollen from species within the same genus clustered closer together in feature space than pollen from more distantly related species. This supports our conclusion that the learned morphological features of grass pollen captured biologically meaningful, taxonomically structured variation and were not likely overfit to the class-specific patterns of individual plants.

### Peak glacial cooling reduced grass diversity on Mt. Kenya

Shannon entropy, estimated from CNN-derived features, provides a proxy for grass diversity without relying on taxonomic identifications (SI Appendix, Fig. S3). Applied to the fossil pollen record from Lake Rutundu, it shows that grass diversity around 3000 m elevation on Mt. Kenya was substantially lower than today during the period of most severe cooling during the LGM and earliest late-Glacial between 21,500 and 16,000 years ago (38, 51), and relatively stable at more or less their modern-day diversity throughout the Holocene (Fig. 3A). Notably, the magnitude of entropy-inferred diversity loss under peak LGM cooling (MAT 7–8 °C below its modern value) is quantitatively consistent with the low entropy values (low species richness) of modern-day grasslands in the upper alpine zone of Mt. Kenya *ca.* 1000 m above Lake Rutundu (ref. 50; Fig. S4), where MAT is ∼6 °C lower (38). This local decrease in grass diversity is likely the result of the depression of the snowline (the 0° C MAT isotherm) and movement of underlying vegetation zones to lower altitudes during Quaternary glacial episodes (45, 46), so that the species-rich heather-moorland grassland around Lake Rutundu was replaced by a species-poor alpine grassland during cooler periods. Additionally, the relatively abrupt drop in entropy (inferred grass diversity) at the onset of peak LGM conditions *ca.* 21,500 years ago, and similarly rapid recovery *ca.* 16,000 years ago (Fig. 3A) suggest a threshold response of Mt. Kenya grasslands involving reduced local viability of many grasses when glacial cooling exceeded >6 °C relative to present-day MAT. Across the PC1 and PC1-PC2 subspaces of CNN-inferred morphological variation, changes in entropy through time are positively correlated with shifts in atmospheric CO_2_ and temperature (Fig. 3C and SI Appendix, Fig. S5), indicating that elevated CO_2_ levels and/or warmer habitat conditions favor more diverse grass communities on Mt. Kenya. This is consistent with multiple studies that have documented higher biomass productivity and biodiversity, particularly in grasslands, under conditions of elevated interglacial CO_2_ and temperature compared to glacial periods (55–58).

The total organic carbon (TOC) concentration, carbon-nitrogen (C/N) ratio and terrestrial biomarker fluxes of Lake Rutundu sediments were low during the LGM before rapidly increasing at *ca.* 16,000 cal. yr BP, indicating a reduction of terrestrial primary productivity in the present-day heathland-moorland zone of Mt. Kenya during the LGM (37, 42, 38). Thus, despite grasses becoming more abundant relative to other herbaceous plants such as heather (Ericaceae), grass productivity and biomass decreased relative to today, because Lake Rutundu was surrounded by the grass-dominated but low-productivity ecosystem now occurring in the upper alpine zone (37, 38, 42). Finally, the positive relationship between temporal variation in entropy across the PC1-PC2 subspace (Fig. 3A) and local biomass burning (as represented by micro-and macrocharcoal influxes; Fig. 3D and SI Appendix, Fig. S5) suggests that climate-driven changes in fire regime may have played an important role in shaping grassland diversity on Mt. Kenya. However, considering that this relationship lacks statistical significance across the other two subspaces of morphological variation, as well as the complex interplay between temperature and moisture effects on fire dynamics in tropical Africa (59), we refrain from deriving strong conclusions from this observation at this time.

### C_4_ grasses have thinner pollen walls and simpler surface texture

The PC1 of CNN features representing pollen morphological variation among modern grass species is significantly associated with whether those grasses use the C_3_ or C_4_ photosynthetic pathway, even after accounting for phylogenetic relatedness (Fig. 5D), indicating that this portion of CNN features captures real morphological differences independent of shared evolutionary history. We cannot directly link PC1 to specific morphological differences between C_3_ and C_4_ grass pollen, but there are statistically significant differences in median pixel brightness and in the entropy of the distribution of pixel values, which may indicate a thicker exine and more complex exine patterning in C_3_ versus C_4_ species (Fig. 5A-C). Morphological differences between the pollen of C_3_ and C_4_ grasses may be linked to their distinct physiology. C_4_ grasses invest more in their root systems and tend to achieve faster growth by producing less dense leaves (higher leaf area to mass ratio), enabling the production of more leaves at the same carbon cost (60). This systematic difference in resource allocation may explain why physiological differences are encoded in pollen morphology, independent of phylogenetic relatedness. We hypothesize that the thinner exine wall and less complex surface patterning of C_4_ grass pollen reflect lower investment in pollen structure resulting from the evolutionary trade-off between resource allocation to roots and shoots.

### Changing prevalence of C_3_ and C_4_ grasses in East African grasslands

During the late Miocene, fire activity was a key factor in the establishment and expansion of C_4_-dominated grasslands in Africa (11, 46, 47), and many contemporary C_4_ grasslands are fire-dependent (61, 63). Our results suggest a similar dynamic over the last 25,000 years on Mt. Kenya. However, we did not find a significant correlation between changes through time in the proportion of C_4_ grasses (Fig. 3B) and precipitation (Fig. 3C and SI Appendix, Fig. S5). Regional records, however, indicate that biomass burning increased during wetter but more seasonal climates in eastern equatorial Africa (∼14,300–12,300 and 11,400–9,600 cal. yr BP), as shown by depleted leaf-wax δD, depleted δ^13^C, and elevated lake levels that mark wetter monsoon phases (64). These wet-dry oscillations may have promoted greater vegetation growth followed by burning, leading to greater C_4_ abundance.

Full-glacial atmospheric CO_2_ concentrations (∼180–200 ppm) are limiting for both C_3_ and C_4_ grasses, but C_4_ species are generally better adapted to lower CO_2_ and drier conditions due to their ability to concentrate CO_2_ around the carboxylating enzyme Rubisco (10) and use water more efficiently (65). However, unlike diversity, the reconstructed proportion of C_4_ grasses correlated only weakly with CO_2_ (SI Appendix, Fig. S5). Likewise, temperature does not appear to have influenced the C_3_:C_4_ ratio through time, with a near-zero correlation. Although C_4_ grasses may have first evolved in warm, tropical environments, they subsequently expanded and diversified into cooler and even cold environments, and today occur across both warm and cold regions (66), notably including cold-tolerant panicoid C_4_ species in modern alpine grasslands (50, 65).

### Implications for deep-time paleoenvironmental reconstruction

Our results demonstrate that deep learning methods applied to superresolution microscopic images can be a powerful tool for revealing shifts in grass diversity and community composition through time, by allowing us to quantify and exploit heretofore hidden morphological variation in fossil grass pollen. By documenting the long-term morphological variability contained in grass pollen assemblages, we can test hypotheses about the evolution and physiological adaptations of grasses over thousands to millions of years and provide a deeper understanding of how grasses adapted to changing climates and environments. The first plants attributed to the Poaceae appeared in the Early Cretaceous (113–101 Mya) (46–48), and by the middle Eocene (∼38 Mya; ref. 4), grasslands were present on every continent except Antarctica. Our study focused on a fossil record of Late Quaternary grasslands to allow validation of the method through direct comparison with other indicators of grass diversity and C_3_:C_4_ ratio. Applying this same approach to fossil grass pollen assemblages further back in the geologic record may at last open a true window on the origin and evolution of this important global biome.

## Methods

### Pollen samples

Pollen samples of 60 extant grass species were isolated from herbarium material from the Missouri Botanical Gardens (USA), University of Illinois (USA), and Harvard (USA) or purchased from Sigma-Aldrich (St. Louis, MO, USA) (SI Appendix, Table S1). These are predominately, but not exclusively, widely distributed species that occur in East Africa, including the important African cereals sorghum (*Sorghum halepense*), finger millet (*Eleusine coracana*), and maize (*Zea mays*) (SI Appendix, Table S2). Over half of our species (56.67%) and over two-thirds of our genera (71.11%) are found on Mt. Kenya. They represent 26.6% of Mt. Kenya’s 124 grass species and 52.6% of its 52 genera (70) (SI Appendix, Table S2). Thirty-six species in our reference pollen dataset are C_3_ grasses and 24 are C_4_ (SI Appendix, Table S1). All but three of the extant grass species were represented by 19 or more pollen grains in our image dataset (SI Appendix, Table S1).

Twenty-four samples of fossil pollen (SI Appendix, Table S4) were isolated from a 7.55-m sediment sequence collected from Lake Rutundu in 1996 (37). Rutundu is a 40-ha oligotrophic lake, 3078 m a.s.l. on the northeast side of Mt. Kenya, just above the current treeline (Fig. 2) (37). Recently deposited pollen was extracted from the surface sediments of Lake Rutundu and five additional small lakes on Mt. Kenya, distributed along an altitudinal transect between 1820 and 4585 m a.s.l. (Fig. 2 and SI Appendix, Table S3), and used as an independent validation dataset (48). All pollen samples were prepared following (71, 72) (details provided in the SI Appendix). Forty-nine to 50 fossil grass grains were imaged from the six Mt. Kenya lake surface sediment samples (SI Appendix, Table S3) and 48–51 fossil grass grains were imaged from each of the 24 Rutundu samples (SI Appendix, Table S4). The ages of the Rutundu samples are derived from an age model based on AMS ^14^C dating of bulk organic carbon (37, 38).

### Paleoenvironmental data

Paleoenvironmental data for the subalpine zone of Mt. Kenya were taken from published studies on the Rutundu sediment record. The temperature reconstruction is based on abundance ratios of preserved glycerol dialkyl glycerol tetraethers (GDGTs) (39) and the precipitation reconstruction is based on the hydrogen-isotope signature of alkanes in fossil leaf waxes from terrestrial vegetation (40). For fire history, we used data on microscopic and macroscopic charcoal influx to Lake Rutundu as a proxy for past variation in local biomass burning and fire prevalence (38). We used the composite record of atmospheric CO_2_ concentration based on air-bubble measurements in several Antarctic ice cores (41).

### SR-SIM imaging

Pollen specimens were imaged using a Zeiss Elyra S1 superresolution structured illumination (SR-SIM) microscope. SR-SIM is an optical microscopy method capable of capturing morphological features <150 nm in length, below the diffraction limit of light (44). We used a 63× Plan Apochromat (1.4 NA) oil objective and an excitation wavelength of 561 nm. Image resolution was 0.0397 µm/pixel. Each axial plane of the Z-stack was constructed with five grid rotations, spaced 0.18 µm apart, with 28–108 planes per grain. Images were cropped to the perimeter of each pollen grain.

### Deep learning architectures and dataset setup

We developed two separate K-way classification CNNs based on the ResNeXt-101 architecture (73): a holistic image CNN (H-CNN) trained on maximum intensity projections (MIP) of entire image stacks, and a patch CNN (P-CNN) trained on square crops of the MIP, each covering ∼10% of the image (33) (Fig. 1A). Both CNNs were trained using images of modern reference pollen (labeled data) and images of fossil pollen (unlabeled data) in a semi-supervised learning (SSL) framework (74). SSL uses the classification model to assign pseudo-labels to unlabeled images and continues to train the model over both the pseudo-labeled and the labeled images. From the pseudo-labeled images, only those that had pseudo-label confidence scores greater ≥95% were retained for training. In our work, we divided the labeled reference data into training (70%) and validation (30%) sets and evaluated model performance using 5-fold cross-validation. The validation set was used for hyperparameter tuning and model selection (SI Appendix, Fig. S1). SSL improved the model’s ability to generalize across both modern and fossil data. Training details are included in the SI Appendix.

The H-CNN generated a single *K*-dimensional logit vector for each pollen specimen, while the P-CNN produced multiple logit vectors corresponding to multiple input patches per specimen (Fig. 1A). The mean of these vectors by P-CNN was calculated and normalized using softmax (33). For each specimen, we fused the outputs of the two CNNs by multiplying the two *K*-dimensional classification probability vectors and normalizing it to unit length (33).

### Feature extraction

Our analysis relied on the features characterized by the two CNNs. We concatenated the 2048-dimensional feature vectors derived from the penultimate (global pooling) layer of the H-CNN and P-CNN to produce a 4096-dimensional feature vector for each specimen (Fig. 1A). This combined feature vector represented a high-level abstraction of the pollen image that served as input for our morphological analyses of diversity and physiology. To visualize and compare morphological variation between the modern grass pollen dataset and the 24 fossil pollen assemblages, we performed PCA on the standardized feature matrix of all modern grains and then mapped fossil grains onto this PCA space (Fig. 4).

### Variation within species and genera

A common assumption in palynology is that variation in pollen from a single individual captures the full range of variation in the morphology of an entire species. To test whether this assumption holds in our dataset and if CNN features captured individual or species-level variability, we conducted two analyses. First, we evaluated whether species from the same genus are closer in feature space than they are to species of different genera. We grouped all modern specimens by species and averaged their CNN feature vectors to obtain species-level embeddings. We then computed pairwise Euclidean distances among all species and used a one-sided Wilcoxon rank-sum test to compare distances within versus between genera. This analysis demonstrated whether there was taxonomic structure in our features. Second, using species in our modern dataset for which pollen samples were available from multiple individuals, as identified in the pollen metadata as originating from MilliporeSigma (Burlington, MA, USA) or “other” sources, we assessed whether variability in feature space differed between pollen samples derived from a single plant and those derived from multiple individuals of the same species. For each species, we measured the average feature variance across specimens, then compared the resulting distributions between the two groups using a Wilcoxon rank-sum test. This analysis demonstrated whether the variability in pollen from a single plant was representative of the diversity within the broader species.

### Ecological simulations and morphological measures of diversity

We first generated artificial communities to test our hypothesis that the morphological diversity within a grass pollen assemblage can serve as a proxy for taxonomic (species) diversity. To avoid reusing the same training data, we randomly withheld half of the modern dataset. Specifically, 30 of the 60 grass species were used to train a new CNN model, while the remaining 30 species (572 specimens) were reserved as an independent test set. This second model was trained solely for the purpose of the diversity-entropy analysis.

In our simulation, each artificial community contained a fixed number of specimens (𝑁 = 50), matching the counts available in fossil assemblages. For each replicate, we first drew species richness

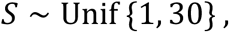

and then sampled 𝑆 species without replacement from the 30-species test pool. Relative abundances were generated as

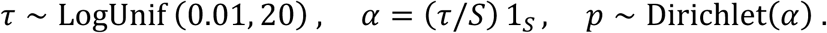

This parametrization allows assemblages to vary from highly uneven (small 𝜏) to highly even (large 𝜏).

Specimen counts were then sampled from a multinomial distribution,

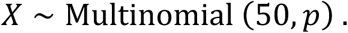

If the sampled count for a species exceeded its available specimens, we capped it at the maximum available and redistributed the excess proportionally among the remaining selected species with remaining availability.

We quantified the diversity of each simulated community using Shannon’s index from the empirical relative frequencies derived from the sampled counts:

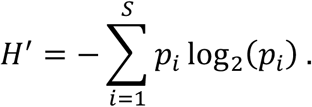

Where 𝑆 is the total number of species and 𝑝_*i*_ is the relative frequency of species 𝑖.

Morphological disparity was quantified from the CNN feature vectors of each assemblage. Feature matrices were standardized and reduced by PCA; each specimen was represented by its PCA scores

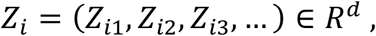

where 𝑑 is the number of retained principal components (PCs). The set of all specimens in a community is: 𝑍 = {𝑍_1_, 𝑍_2_, …, 𝑍_*n*_} ⊂ 𝑅^*d*^, where 𝑛 is the number of specimens in an assemblage.

We estimated the continuous analogue of Shannon entropy, the differential entropy of the score distribution. For a density function 𝑓(𝑥), differential entropy is defined as

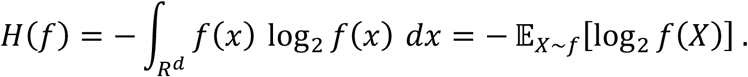

This measures uncertainty associated with drawing a specimen at random from the morphological space. Because 𝑓 is unknown, we estimated it directly from the observed specimens in each assemblage using Gaussian kernel density estimation (KDE) based only on the observed PCA scores:

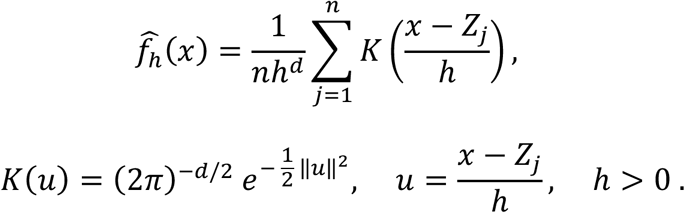

Using a leave-one-out (LOO) version to reduce self-influence:

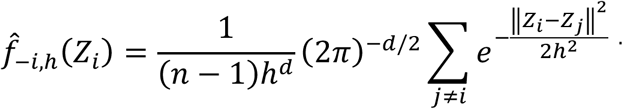

Since 𝐻(𝑓) = −𝔼[log_2_ 𝑓 (𝑋)], we estimate entropy by approximating the expectation with the empirical mean:

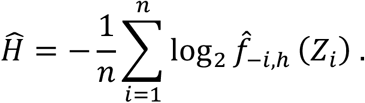

Bandwidths were selected by maximizing the mean LOO log-likelihood of the pooled PCA scores, separately for 𝑑 ∈ {1, 2, 3}, evaluated over 500 logarithmically spaced values between 0.01 and 10. The optimum for each 𝑑 was denoted as ℎ^∗^. For the Rutundu fossil assemblages, entropy was computed only at ℎ^∗^. For the simulated assemblages and surface sediment samples, we evaluated sensitivity by scaling ℎ^∗^ in log_2_ space:

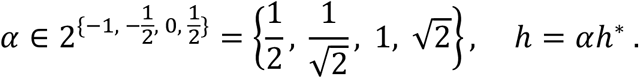

These four values bracket the optimal ℎ^∗^ and allow evaluation of how smoothing influences diversity reconstruction.

Finally, morphological disparity from the simulated communities was compared with their Shannon diversity values across 100 replicates using linear regression, with Pearson’s correlation coefficient (r) and its significance reported. For each panel, we also display uncertainty as 95% confidence bands for the mean response and 95% prediction bands for new observations (SI Appendix, Fig. S3).

### Surface sample analysis

We applied the entropy measure described above to pollen isolated from the surface sediments of six Mt. Kenya lakes, including Lake Rutundu. For each site, Shannon entropy (bits) was calculated from PCA scores of the pollen assemblage and compared with Poaceae species richness measured in vegetation surveys along the same elevational transect (50). Analyses were performed on three PCA subspaces (PC1, PC1-PC2, and PC1-PC2-PC3). To test the robustness of the entropy estimates, we repeated the analysis across the same four bandwidth scaling factors defined above. To visualize elevational diversity patterns, site-level entropy values were connected using a piecewise cubic Hermite interpolating polynomial (PCHIP) implemented in SciPy (v. 1.16.0) (63).

### Fossil grass diversity estimation and paleoenvironmental correlates

We next applied the same entropy measure to 24 fossil pollen assemblages isolated from the Lake Rutundu sediment core, spanning 25,000 years. For each assemblage, we calculated Shannon entropy across the PC1, PC1-PC2, and PC1-PC2-PC3 subspaces. These diversity estimates were then compared with five paleoenvironmental variables: atmospheric CO_2_ level, local reconstructed temperature and precipitation, and micro- and macrocharcoal influx as proxy for fire prevalence.

Correlations were evaluated using Pearson’s coefficient (r) and associated p-values (SI Appendix, Fig. S5). All environmental proxy series were smoothed with a 4-point moving average prior to visualization and correlation analysis to reduce high-frequency noise.

### Grass pollen morphology and photosynthetic pathway

We averaged the CNN feature matrix by species and performed a standard PCA. We tested whether the principal components discriminated between C_3_ and C_4_ photosynthetic pathways using Welch’s t-tests (Fig. 5).

To determine whether the observed differences reflect genuine functional differentiation or phylogenetic history, we tested whether the principal components remained significantly discriminatory when accounting for evolutionary relationships. We placed all 60 grass species within a recent time-calibrated Poaceae phylogeny of ∼90% of extant grass genera (69) by creating polytomies of multiple congeneric species using the function bind.tip in the R package ape (76), with branch lengths of 0.001. This conservative approach inflated phylogenetic signal (77), but allowed us to include all species in the analysis. We tested for phylogenetic signal in PC scores using Pagel’s λ with the phylosig function in the R package phytools (78), then applied phylogenetic generalized least squares (PGLS) regression using the phylolm function in the R package phylolm (79) to determine whether C3–C4 differences persisted after accounting for shared ancestry. We set all branch lengths to equal values to address potential biases from artificial polytomies.

### Estimates of past C_4_ abundance and paleoenvironmental correlates

We performed a principal component analysis (PCA) on the CNN features of all individual pollen specimens from our modern Poaceae dataset (766 C_3_ and 475 C_4_ grains), retaining the principal components explaining up to 95% of the variance. The fossil pollen CNN feature matrix was standardized using the mean and standard deviation of the modern data and projected onto this PCA space. We then trained a gradient-boosted decision tree classifier (26, 27), implemented in the XGBoost Python library (v. 2.1.4) (80), and evaluated it with stratified five-fold cross-validation. The final trained model was applied to each fossil specimen from the 24 Rutundu samples to infer its photosynthetic pathway (C_3_ or C_4_) (Fig. 1C). For comparison, we also report in the same plot the δ^13^C-based Bayesian mixture model estimates of C_4_ abundances (38) and the fossil cuticle identifications of C_4_ taxa (37), each shown as originally reported in their respective age models (Fig. 3B). For the latter, we traced the upper and lower bounding curves from their published figure and linearly interpolated each onto a dense evenly spaced age grid (≤25 Ka). The midpoint (arithmetic mean) of the two bounds was taken as the central trajectory, with the half-difference forming an uncertainty envelope. We show shaded intervals representing 95% Wilson confidence intervals (81) for our estimates, 95% credible intervals for the Bayesian mixture model (38), and the range between “certain only” versus “certain + likely” cuticle identifications (37).

To evaluate temporal consistency between C_4_ records, all series were smoothed using cubic smoothing splines (82) with varying smoothing factors (*α* = 0.01–0.4), interpolated onto a common 1000-point age grid spanning the 25 ka BP–present interval, and compared pairwise using Pearson’s correlation coefficient (r) (SI Appendix, Fig. S6). We also assessed the correlation between our time series of estimated C_4_ abundance and those of the five paleoenvironmental variables: atmospheric CO_2_ concentration, local temperature, local precipitation, and micro- and macro-charcoal influx. We calculated Pearson’s r and its corresponding p-value for each individual regression (SI Appendix, Fig. S5).

### Quantifying morphological differences between C_3_ and C_4_ pollen

We used entropy to estimate the complexity of the grass pollen surface texture, with higher values indicating more random and complex textural patterning. We calculated Shannon entropy by segmenting the pollen grain image and quantifying the isolated grain’s information content (complexity) based on the distribution of its pixel intensities (83). Because pollen exine is autofluorescent, pixel intensity allows us to infer variations in thickness of the pollen wall, with brighter images suggesting a thicker wall (44). We used direct exine thickness measurements (24) for 35 C_3_ and 23 C_4_ grass species to confirm this relationship. We calculated species-level medians of pixel intensity, morphological complexity (measured using Shannon entropy), and exine thickness across all specimens of each species and used Welch’s t-test to assess the statistical significance of the observed differences between C_3_ and C_4_ species (Fig. 5).

## Acknowledgments

Imaging costs and postdoctoral support for M.A.U. were provided by NSF-DBI – Advances in Bioinformatics (NSF-DBI-1262561) to S.W.P. S.K. was supported in part by the University of Macau (SRG2023-00044-FST), Science and Technology Development Fund of Macau SAR (0067/2024/ITP2), and the Institute of Collaborative Innovation. M.-E.A. was supported in part by the University of Illinois Tom L. Phillips Memorial Fund for Paleobotany. We thank Tim Gallaher for referring us to his recently published grass phylogeny.

## Author Contributions

M.-E.A., S.K., and S.W.P. designed the research. M.-E.A. conducted the computational analyses and experiments. M.A.U. conducted the imaging, sample preparation, and lab analyses. F.A.S.-P. and D.V. provided samples and collected data. S.W.P. and S.K. supervised the research. M.-E.A., S.W.P., S.K., and M.A.U. wrote the manuscript, with feedback from all authors.

## Data Availability

Superresolution structured illumination images used in this study were submitted to the Illinois Databank (DOI: 10.13012/B2IDB-1391373_V1) (84). The code base developed for this study is publicly available on GitHub (85) and can be accessed using the following link: https://github.com/paleopollen/Pollen_Diversity_Dynamics. In addition, several components of the analytical pipeline were adapted from code published in (86) available at: https://github.com/paleopollen/Novel_Pollen_Phylogenetic_Placement.

