## Supplementary Information for "Deep learning of fossil pollen morphology reveals 25,000 years of ecological change in East African grasslands"

**This PDF file includes:**

Supporting text
Figures S1 to S6
Tables S1 to S5
SI References

**Supporting Information Text**

***Study site***

Our study site is Mt. Kenya, an extinct volcano located east of the Kenyan Rift Valley in eastern equatorial Africa. This region experiences a bimodal pattern of seasonal rainfall with ‘‘long’’ rains from March to May and “short” rains from October to December. As most rainfall is sourced in the Indian Ocean, precipitation is heaviest (>2500 mm/yr) at mid-elevations (2000–3000 m) on the southeastern flank and lightest (<1500 mm/yr) on the northern flank (1, 2). Above the treeline, at ~2900–3400 m a.s.l., total precipitation declines with increased elevation to <900 mm/yr at 4500 m.

Precipitation and temperature dynamics on Mt. Kenya over the past 25,000 years are well understood from previous research on the sediment records of local high-elevation lakes, and include reconstructions of past moisture balance (3, 4), precipitation (5), and temperature (6, 7). Air temperatures during the Last Glacial Maximum (LGM) were 3–4.5°C lower than today, with most warming occurring during the transition into the Holocene (6). Precipitation also decreased during the LGM by 25–35% on average due to weaker monsoonal moisture advection associated with lower sea surface temperatures (SST) in the Indian Ocean (3, 4, 8).

Vegetation on present-day Mt. Kenya is delineated into distinct altitudinal zones (Fig. 2). The base of the mountain is surrounded by $C_4$-grass-dominated savanna and cultivated areas. The savanna transitions upwards into a montane rainforest on all but the northern slope, which is dominated by a wooded grassland with *Acacia* and *Euphorbia*. Above the treeline, vegetation first grades into a subalpine Ericaceous belt characterized by tall, woody, microphyllous shrubs (*Erica*, *Anthospermum*, and *Philippia*) with an understory of $C_4$ and $C_3$ grasses; above that the Afroalpine Belt, characterized by $C_3$ tussock grassland; and finally the Nival Zone of mostly bare rock and ice (9–11).

In addition to the 25,000-year sediment record of Lake Rutundu, recently deposited surface sediments were collected from five additional small (<51 ha) lakes distributed along an altitudinal transect between 1820 and 4585 m on the northeast flank of Mt. Kenya (Fig. 2). Lakes Rutundu and Nkunga are volcanic maar lakes, whereas Lake Ellis, Small Hall Tarn, and Simba Tarn occupy small ice-scoured basins. Lake Nkunga is located in the montane rainforest and is surrounded by sedges, ferns and aquatic $C_4$ grasses. Lakes Rutundu and Ellis are located above the treeline in the Ericaceous belt. Small Hall Tarn is located in an Afroalpine $C_3$ shrub grassland, while Simba Tarn lies close to the year-round freezing level where plant growth is extremely limited (9).

***Modern grass pollen samples***

Modern grass pollen samples were procured from within mostly closed flowers of herbarium specimens stored at the Missouri Botanical Gardens Herbarium (USA), University of Illinois Herbarium (USA), and Harvard Herbarium (USA). *Dactylis glomerata*, *Secale cereale*, and *Sorghum halepense* pollen were obtained from Sigma-Aldrich (St. Louis, MO, USA). For each modern sample, 3 to 37 pollen grains, typically ~20, were obtained from two to four flowers (Table S1). Only 3 and 4 pollen grains could be obtained from, respectively, *Andropogon chrysostachyus* and *Eleusine coracana*. Harvard Herbarium and Sigma-Aldrich samples were prepared using a modified standard procedure (12) that excluded carbon-containing compounds (13, 14). Samples from the Missouri Botanical Gardens and University of Illinois were prepared following (15).

***Fossil grass pollen samples***

Samples of fossil grass pollen were obtained from a 7.55 m sediment sequence recovered from the central area of Lake Rutundu in 1996 and from recently deposited surface sediments in the five other lakes. These samples were originally used for carbon-isotope analysis of individual grass pollen grains following the preparation protocol described in (13, 16). Briefly, we extracted grass pollen from ~1 $cm^3$ sediment samples taken at 24 depth intervals in the Rutundu sediment core, and processed them following standard techniques (12) with modifications to exclude carbon-containing compounds (17).

##### *Staining and mounting of pollen samples*

Sporopollenin, the biopolymer forming a major component of the resistant outer (exine) wall of pollen grains, is naturally autofluorescent. However, fluorescent labeling can increase the signal-to-noise ratio (SNR) and vastly improve the quality of SR-SIM images (18). All pollen samples were washed a minimum of three times with ultrapure water in 10 µm cell strainers. Pollen in the cell strainers was incubated in periodic acid ($HIO_4$, 1 g/dL) for a minimum of 8 hours followed by three washes in distilled water. The pollen was then incubated with Schiff's reagent for a minimum of 30 min or until the pollen began to change color. Pollen was washed to remove excess stain and to allow the color to fully develop.

Stained pollen samples were mounted in Eukitt, a solid resin-based medium with a refractive index close to that of glass, to ensure successful superresolution imaging (18). The pollen was first dehydrated in the 10 µm cell strainers with a graded isopropanol/xylene series of 70%, 80%, 100% isopropanol followed by 2:1, 1:1, 1:2 solutions of ethanol:xylene. After dehydration, pollen was transferred to microcentrifuge tubes and incubated in 100% xylene with 3 drops of Eukitt to allow infiltration of the mounting medium overnight. Excess xylene was then removed and additional Eukitt was added to cover the pollen sample. The samples were stirred to distribute the pollen evenly within the mounting medium. A 500 µL pipette was then used to place a drop of about 3–4 mm in diameter on a glass microscope slide. The sample drop was covered with a high-performance cover glass (0.17 mm thickness). The mounted samples were dried overnight at room temperature in a laboratory hood.

##### *Computational neural network training*

We applied data augmentation to both the labeled modern and the unlabeled fossil pollen image datasets to enhance the robustness and generalization capability of the CNN models. Weak augmentation methods, including random vertical and horizontal flips and rotations within the range of [-90°, +90°], were applied to each specimen with a probability of 0.5. For the unlabeled fossil pollen, additional strong augmentation techniques were used to encourage the model to produce consistent outputs for different strongly distorted versions of the same input image, thus further improving generalization. These techniques included random sharpness adjustments, auto contrast, and equalization (i.e., adjusting the sharpness, maximizing the contrast, and equalizing the histogram of a given image randomly, respectively), each applied with a probability of 0.5. The combination of these standard augmentation strategies is based on functions provided by the PyTorch library (19). All images were resized to $224 \times 224 \times 3$ pixel resolution using bilinear interpolation before being input to the CNN.

*Labeled data learning loss*

The CNN was trained on labeled data by minimizing the categorical cross-entropy loss,

$$L_{CE} = -\frac{1}{N}\sum_{i=1}^{N}\sum_{k=1}^{K} T_{i,k} \log\left(P_{i,k}\right)$$

Here, $N$ is the number of specimens in the labeled dataset, $K$ is the number of classes, $T_{i,k}$ is an indicator variable equal to 1 if specimen $i$ belongs to class $k$ (and 0 otherwise), and $P_{i,k}$ is the predicted probability that specimen $i$ belongs to class $k$ (20).

*Unlabeled data learning loss*

For each unlabeled specimen $j$, the model produces a vector of class probabilities from the weakly augmented image,

$$\hat{y}^{(j)} = \left(\hat{y}_1^{(j)}, \dots, \hat{y}_K^{(j)}\right).$$

The pseudo-label for specimen $j$ is defined as the class index

$$\hat{c}^{(j)} = \arg\max_k \hat{y}_k^{(j)} .$$

Only pseudo-labels whose maximum predicted probability exceeds a confidence threshold $\tau$ are retained. The unlabeled loss is defined as

$$L^u = -\frac{\lambda_u}{M}\sum_{j=1}^{M} \mathbf{1}\left(\max_k \hat{y}_k^{(j)} > \tau\right) \log\left(P_{\hat{c}^{(j)},j}^{\text{strong}}\right) .$$

Here, $M$ is the number of unlabeled specimens, $\lambda_u$ is a balancing factor for the unlabeled loss, and $P_{\hat{c}^{(j)},j}^{\text{strong}}$ denotes the predicted probability assigned to the pseudo-label $\hat{c}^{(j)}$ for specimen $j$, evaluated on the strongly augmented version of the image. The indicator function takes a value of 1 when the maximum predicted class probability exceeds the threshold $\tau$, and 0 otherwise. Only high-confidence pseudo-labeled specimens contribute to the unlabeled loss.

*Total training objective*

The total loss for each training batch is given by the sum of the labeled and unlabeled losses,

$$L = L_{CE} + L^{u}.$$

Optimization was performed using stochastic gradient descent with a momentum of 0.9. The initial learning rate was set to 0.0009 and reduced by a factor of two every two epochs using a learning rate scheduler. Models were trained for 30 epochs with a batch size of 10.

Fig. S1. Confusion matrices depicting CNN classification accuracies for the 60 modern Poaceae species across five training/validation splits and their average. Shown are the H-CNN (top left), P-CNN (top right), and the fused probabilities (bottom). Rows represent the true taxon, with the number of specimens imaged per taxon in parentheses. Columns represent the model’s predictions, with the classification accuracy for each species in parentheses.

### Split 1

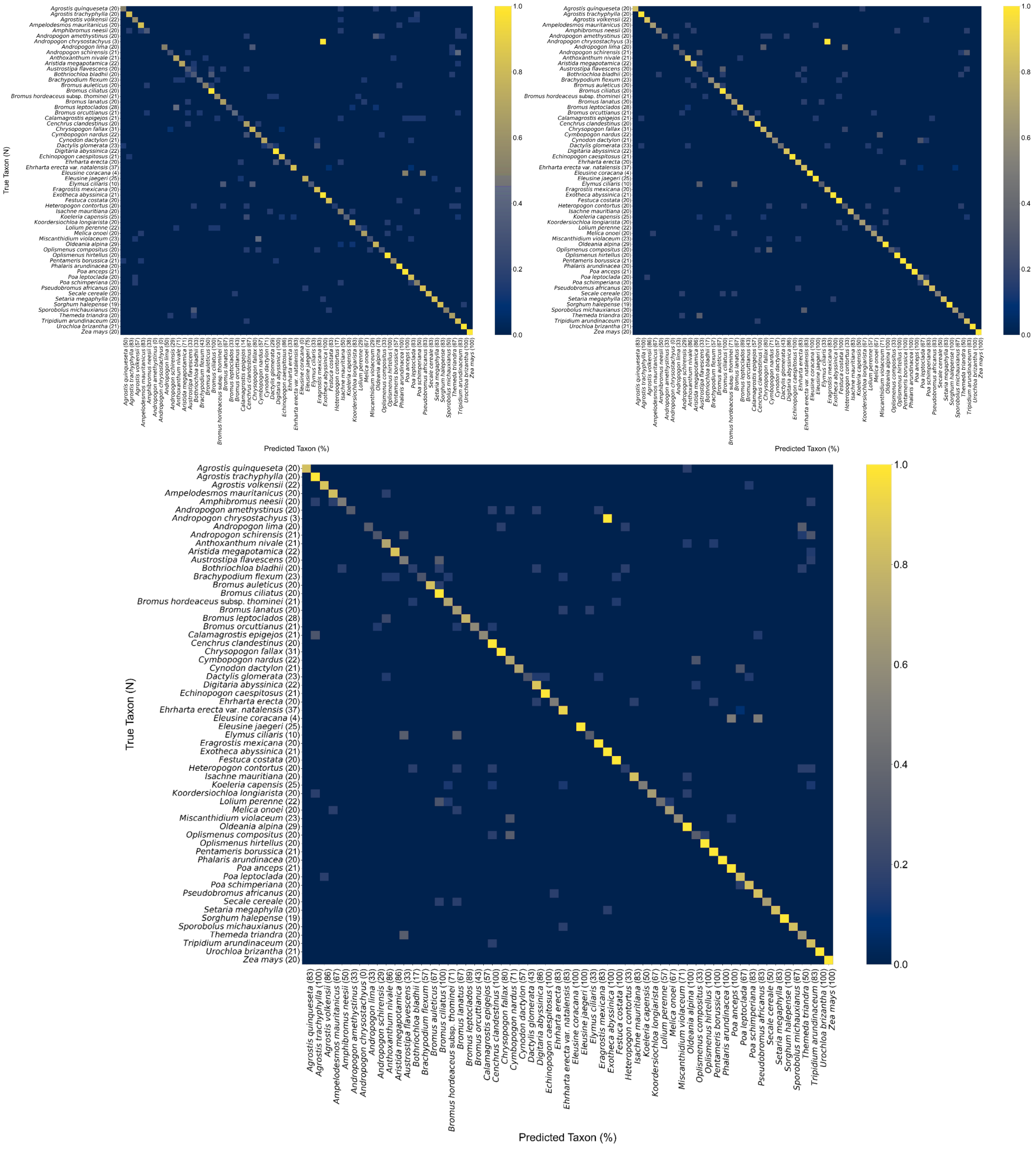
True Taxon (N)
Predicted Taxon (%)
Agrostis quinqueseta (20)
Agrostis trachyphylla (20)
Agrostis volkensii (22)
Ampelodesmos mauritanicus (20)
Amphibromus neesii (20)
Andropogon amethystinus (20)
Andropogon chrysostachyus (3)
Andropogon lima (20)
Andropogon schirensis (21)
Anthoxanthum nivale (21)
Aristida megapotamica (22)
Austrostipa flavescens (20)
Bothriochloa bladhii (20)
Brachypodium flexum (23)
Bromus auleticus (20)
Bromus ciliatus (20)
Bromus hordeaceus subsp. thominei (21)
Bromus lanatus (20)
Bromus leptoclados (28)
Bromus orcuttianus (21)
Calamagrostis epigejos (21)
Cenchrus clandestinus (20)
Chrysopogon fallax (31)
Cymbopogon nardus (22)
Cynodon dactylon (21)
Dactylis glomerata (23)
Digitaria abyssinica (22)
Echinopogon caespitosus (21)
Ehrharta erecta (20)
Ehrharta erecta var. natalensis (37)
Eleusine coracana (4)
Eleusine jaegeri (25)
Elymus ciliaris (10)
Eragrostis mexicana (20)
Exotheca abyssinica (21)
Festuca costata (20)
Heteropogon contortus (20)
Isachne mauritiana (20)
Koeleria capensis (25)
Koordersiochloa longiarista (20)
Lolium perenne (22)
Melica onoei (20)
Miscanthidium violaceum (23)
Oldeania alpina (29)
Oplismenus compositus (20)
Oplismenus hirtellus (20)
Pentameris borussica (21)
Phalaris arundinacea (20)
Poa anceps (21)
Poa leptoclada (20)
Poa schimperiana (20)
Pseudobromus africanus (20)
Secale cereale (20)
Setaria megaphylla (20)
Sorghum halepense (19)
Sporobolus michauxianus (20)
Themeda triandra (20)
Tripidium arundinaceum (20)
Urochloa brizantha (21)
Zea mays (20)
Agrostis quinqueseta (83)
Agrostis trachyphylla (100)
Agrostis volkensii (86)
Ampelodesmos mauritanicus (67)
Amphibromus neesii (50)
Andropogon amethystinus (33)
Andropogon chrysostachyus (0)
Andropogon lima (33)
Andropogon schirensis (29)
Anthoxanthum nivale (86)
Aristida megapotamica (86)
Austrostipa flavescens (33)
Bothriochloa bladhii (17)
Brachypodium flexum (57)
Bromus auleticus (67)
Bromus ciliatus (100)
Bromus hordeaceus subsp. thominei (71)
Bromus lanatus (67)
Bromus leptoclados (89)
Bromus orcuttianus (43)
Calamagrostis epigejos (57)
Cenchrus clandestinus (100)
Chrysopogon fallax (80)
Cymbopogon nardus (71)
Cynodon dactylon (57)
Dactylis glomerata (43)
Digitaria abyssinica (86)
Echinopogon caespitosus (100)
Ehrharta erecta (83)
Ehrharta erecta var. natalensis (83)
Eleusine coracana (100)
Eleusine jaegeri (100)
Elymus ciliaris (33)
Eragrostis mexicana (83)
Exotheca abyssinica (100)
Festuca costata (100)
Heteropogon contortus (33)
Isachne mauritiana (83)
Koeleria capensis (50)
Koordersiochloa longiarista (67)
Lolium perenne (57)
Melica onoei (67)
Miscanthidium violaceum (71)
Oldeania alpina (100)
Oplismenus compositus (33)
Oplismenus hirtellus (100)
Pentameris borussica (100)
Phalaris arundinacea (100)
Poa anceps (100)
Poa leptoclada (67)
Poa schimperiana (83)
Pseudobromus africanus (83)
Secale cereale (50)
Setaria megaphylla (83)
Sorghum halepense (100)
Sporobolus michauxianus (67)
Themeda triandra (50)
Tripidium arundinaceum (83)
Urochloa brizantha (100)
Zea mays (100)
1.0
0.8
0.6
0.4
0.2
0.0

### Split 2

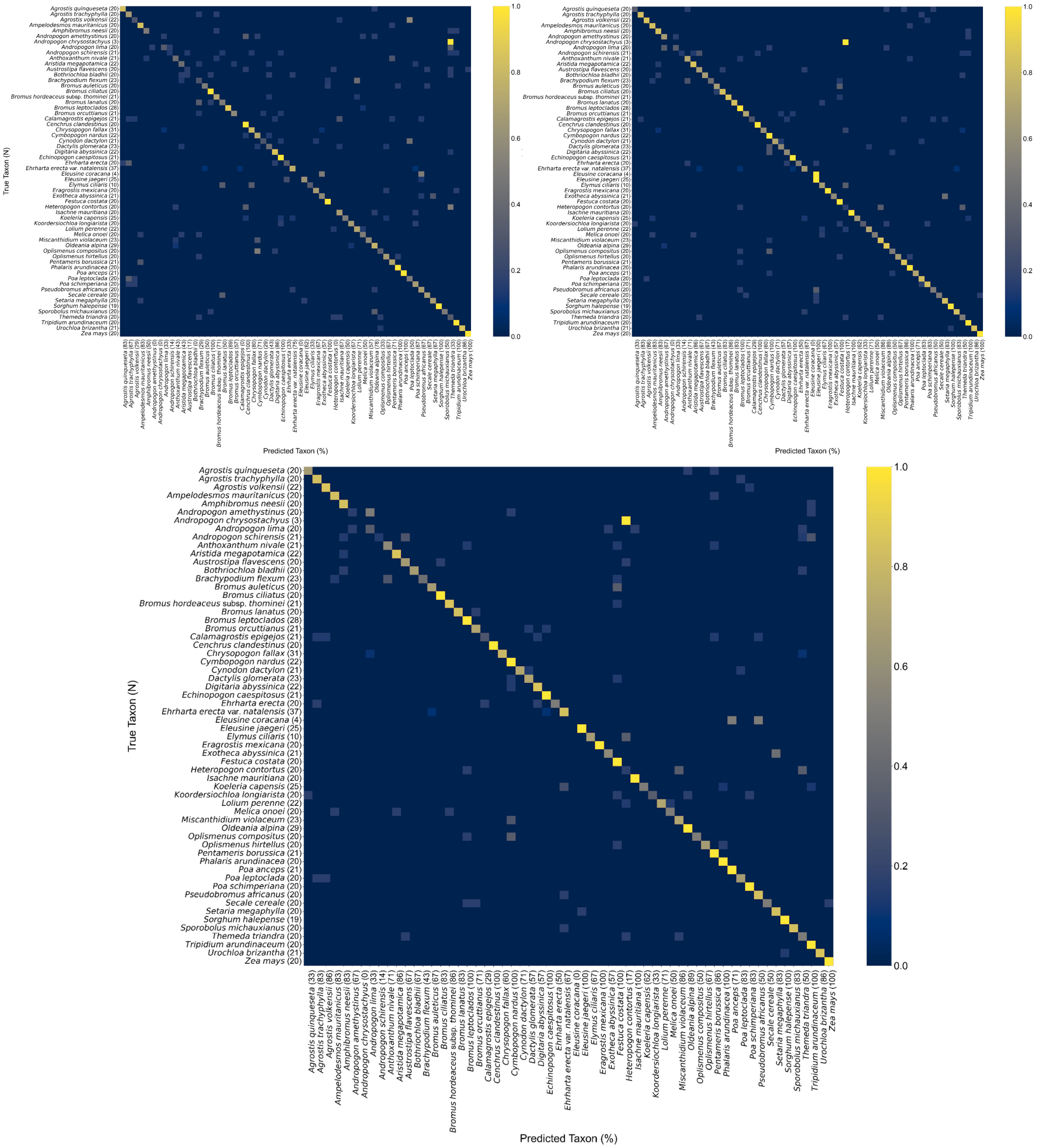
True Taxon (N)
Predicted Taxon (%)
Agrostis quinqueseta (20)
Agrostis trachyphylla (20)
Agrostis volkensii (22)
Ampelodesmos mauritanicus (20)
Amphibromus neesii (20)
Andropogon amethystinus (20)
Andropogon chrysostachyus (3)
Andropogon lima (20)
Andropogon schirensis (21)
Anthoxanthum nivale (21)
Aristida megapotamica (22)
Austrostipa flavescens (20)
Bothriochloa bladhii (20)
Brachypodium flexum (23)
Bromus auleticus (20)
Bromus ciliatus (20)
Bromus hordeaceus subsp. thominei (21)
Bromus lanatus (20)
Bromus leptoclados (28)
Bromus orcuttianus (21)
Calamagrostis epigejos (21)
Cenchrus clandestinus (20)
Chrysopogon fallax (31)
Cymbopogon nardus (22)
Cynodon dactylon (21)
Dactylis glomerata (23)
Digitaria abyssinica (22)
Echinopogon caespitosus (21)
Ehrharta erecta (20)
Ehrharta erecta var. natalensis (37)
Eleusine coracana (4)
Eleusine jaegeri (25)
Elymus ciliaris (10)
Eragrostis mexicana (20)
Exotheca abyssinica (21)
Festuca costata (20)
Heteropogon contortus (20)
Isachne mauritiana (20)
Koeleria capensis (25)
Koordersiochloa longiarista (20)
Lolium perenne (22)
Melica onoei (20)
Miscanthidium violaceum (23)
Oldeania alpina (29)
Oplismenus compositus (20)
Oplismenus hirtellus (20)
Pentameris borussica (21)
Phalaris arundinacea (20)
Poa anceps (21)
Poa leptoclada (20)
Poa schimperiana (20)
Pseudobromus africanus (20)
Secale cereale (20)
Setaria megaphylla (20)
Sorghum halepense (19)
Sporobolus michauxianus (20)
Themeda triandra (20)
Tripidium arundinaceum (20)
Urochloa brizantha (21)
Zea mays (20)
Agrostis quinqueseta (33)
Agrostis trachyphylla (83)
Agrostis volkensii (86)
Ampelodesmos mauritanicus (83)
Amphibromus neesii (83)
Andropogon amethystinus (67)
Andropogon chrysostachyus (0)
Andropogon lima (33)
Andropogon schirensis (14)
Anthoxanthum nivale (71)
Aristida megapotamica (86)
Austrostipa flavescens (67)
Bothriochloa bladhii (67)
Brachypodium flexum (43)
Bromus auleticus (67)
Bromus ciliatus (83)
Bromus hordeaceus subsp. thominei (86)
Bromus lanatus (83)
Bromus leptoclados (100)
Bromus orcuttianus (71)
Calamagrostis epigejos (29)
Cenchrus clandestinus (100)
Chrysopogon fallax (60)
Cymbopogon nardus (100)
Cynodon dactylon (71)
Dactylis glomerata (57)
Digitaria abyssinica (57)
Echinopogon caespitosus (100)
Ehrharta erecta (50)
Ehrharta erecta var. natalensis (67)
Eleusine coracana (0)
Eleusine jaegeri (100)
Elymus ciliaris (67)
Eragrostis mexicana (100)
Exotheca abyssinica (57)
Festuca costata (100)
Heteropogon contortus (17)
Isachne mauritiana (100)
Koeleria capensis (62)
Koordersiochloa longiarista (33)
Lolium perenne (71)
Melica onoei (50)
Miscanthidium violaceum (86)
Oldeania alpina (89)
Oplismenus compositus (50)
Oplismenus hirtellus (50)
Pentameris borussica (86)
Phalaris arundinacea (100)
Poa anceps (71)
Poa leptoclada (83)
Poa schimperiana (83)
Pseudobromus africanus (50)
Secale cereale (50)
Setaria megaphylla (83)
Sorghum halepense (100)
Sporobolus michauxianus (83)
Themeda triandra (50)
Tripidium arundinaceum (100)
Urochloa brizantha (86)
Zea mays (100)
1.0
0.8
0.6
0.4
0.2
0.0

### Split 3

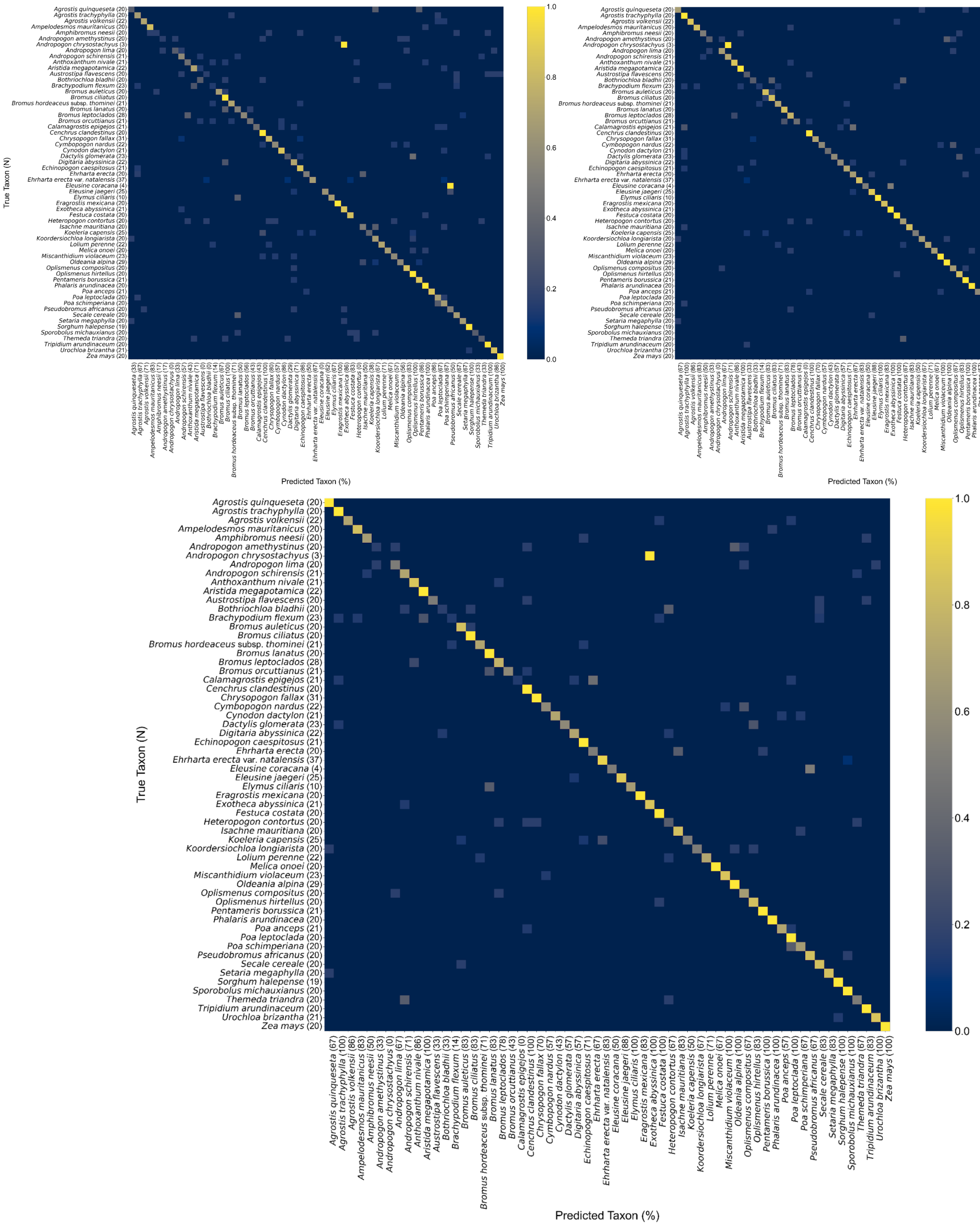

### Split 4

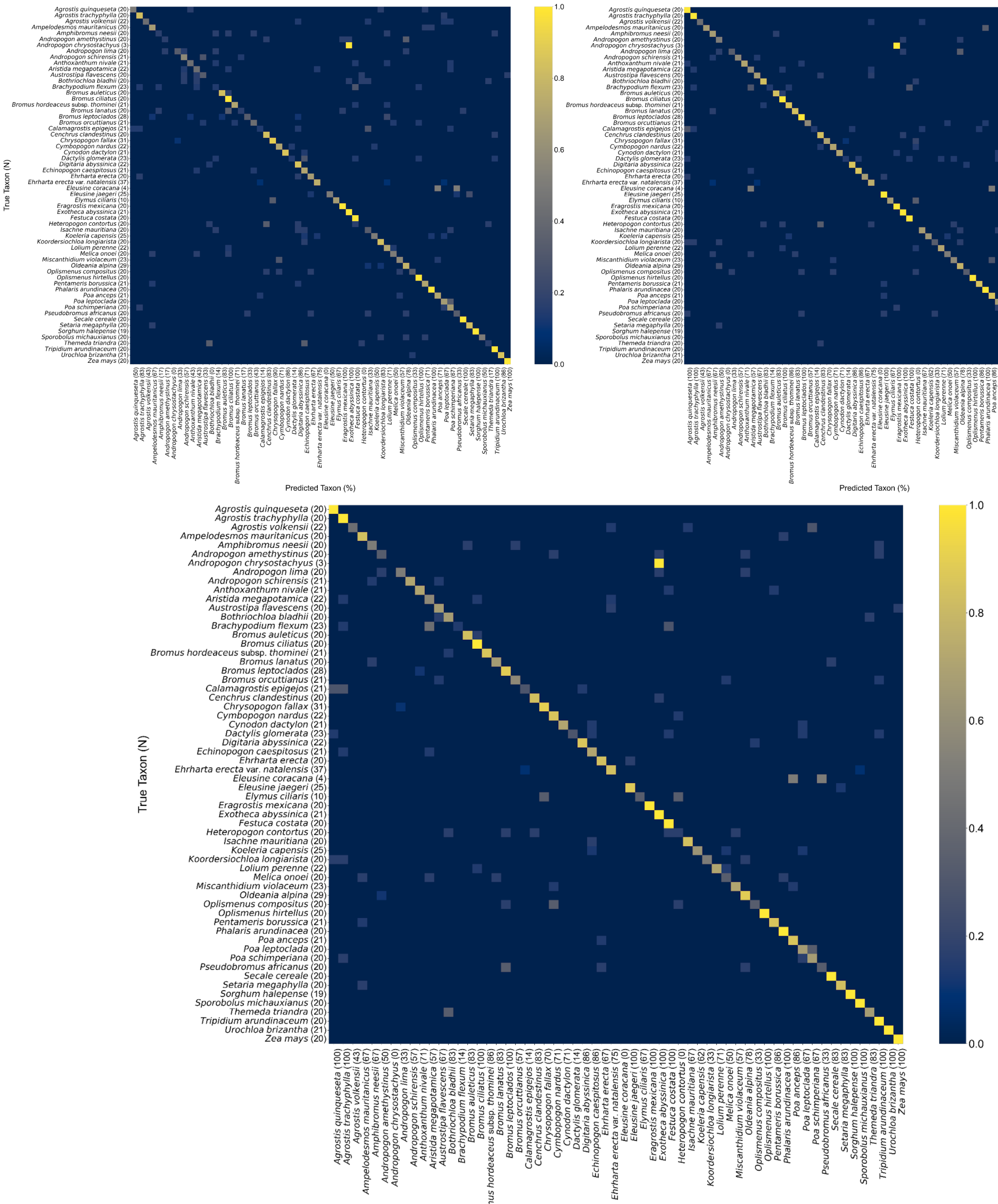
True Taxon (N)
Predicted Taxon (%)
Agrostis quinqueseta (20)
Agrostis trachyphylla (20)
Agrostis volkensii (22)
Ampelodesmos mauritanicus (20)
Amphibromus neesii (20)
Andropogon amethystinus (20)
Andropogon chrysostachyus (3)
Andropogon lima (20)
Andropogon schirensis (21)
Anthoxanthum nivale (21)
Aristida megapotamica (22)
Austrostipa flavescens (20)
Bothriochloa bladhii (20)
Brachypodium flexum (23)
Bromus auleticus (20)
Bromus ciliatus (20)
Bromus hordeaceus subsp. thominei (21)
Bromus lanatus (20)
Bromus leptoclados (28)
Bromus orcuttianus (21)
Calamagrostis epigejos (21)
Cenchrus clandestinus (20)
Chrysopogon fallax (31)
Cymbopogon nardus (22)
Cynodon dactylon (21)
Dactylis glomerata (23)
Digitaria abyssinica (22)
Echinopogon caespitosus (21)
Ehrharta erecta (20)
Ehrharta erecta var. natalensis (37)
Eleusine coracana (4)
Eleusine jaegeri (25)
Elymus ciliaris (10)
Eragrostis mexicana (20)
Exotheca abyssinica (21)
Festuca costata (20)
Heteropogon contortus (20)
Isachne mauritiana (20)
Koeleria capensis (25)
Koordersiochloa longiarista (20)
Lolium perenne (22)
Melica onoei (20)
Miscanthidium violaceum (23)
Oldeania alpina (29)
Oplismenus compositus (20)
Oplismenus hirtellus (20)
Pentameris borussica (21)
Phalaris arundinacea (20)
Poa anceps (21)
Poa leptoclada (20)
Poa schimperiana (20)
Pseudobromus africanus (20)
Secale cereale (20)
Setaria megaphylla (20)
Sorghum halepense (19)
Sporobolus michauxianus (20)
Themeda triandra (20)
Tripidium arundinaceum (20)
Urochloa brizantha (21)
Zea mays (20)
Agrostis quinqueseta (100)
Agrostis trachyphylla (100)
Agrostis volkensii (43)
Ampelodesmos mauritanicus (67)
Amphibromus neesii (67)
Andropogon amethystinus (50)
Andropogon chrysostachyus (0)
Andropogon lima (33)
Andropogon schirensis (57)
Anthoxanthum nivale (71)
Aristida megapotamica (43)
Austrostipa flavescens (67)
Bothriochloa bladhii (83)
Brachypodium flexum (14)
Bromus auleticus (83)
Bromus ciliatus (100)
Bromus hordeaceus subsp. thominei (86)
Bromus lanatus (83)
Bromus leptoclados (100)
Bromus orcuttianus (57)
Calamagrostis epigejos (14)
Cenchrus clandestinus (83)
Chrysopogon fallax (70)
Cymbopogon nardus (71)
Cynodon dactylon (71)
Dactylis glomerata (14)
Digitaria abyssinica (86)
Echinopogon caespitosus (86)
Ehrharta erecta (67)
Ehrharta erecta var. natalensis (75)
Eleusine coracana (0)
Eleusine jaegeri (100)
Elymus ciliaris (67)
Eragrostis mexicana (100)
Exotheca abyssinica (100)
Festuca costata (100)
Heteropogon contortus (0)
Isachne mauritiana (67)
Koeleria capensis (62)
Koordersiochloa longiarista (33)
Lolium perenne (71)
Melica onoei (50)
Miscanthidium violaceum (57)
Oldeania alpina (78)
Oplismenus compositus (33)
Oplismenus hirtellus (100)
Pentameris borussica (86)
Phalaris arundinacea (100)
Poa anceps (86)
Poa leptoclada (67)
Poa schimperiana (67)
Pseudobromus africanus (33)
Secale cereale (83)
Setaria megaphylla (83)
Sorghum halepense (100)
Sporobolus michauxianus (100)
Themeda triandra (83)
Tripidium arundinaceum (100)
Urochloa brizantha (100)
Zea mays (100)
0.0
0.2
0.4
0.6
0.8
1.0

### Split 5

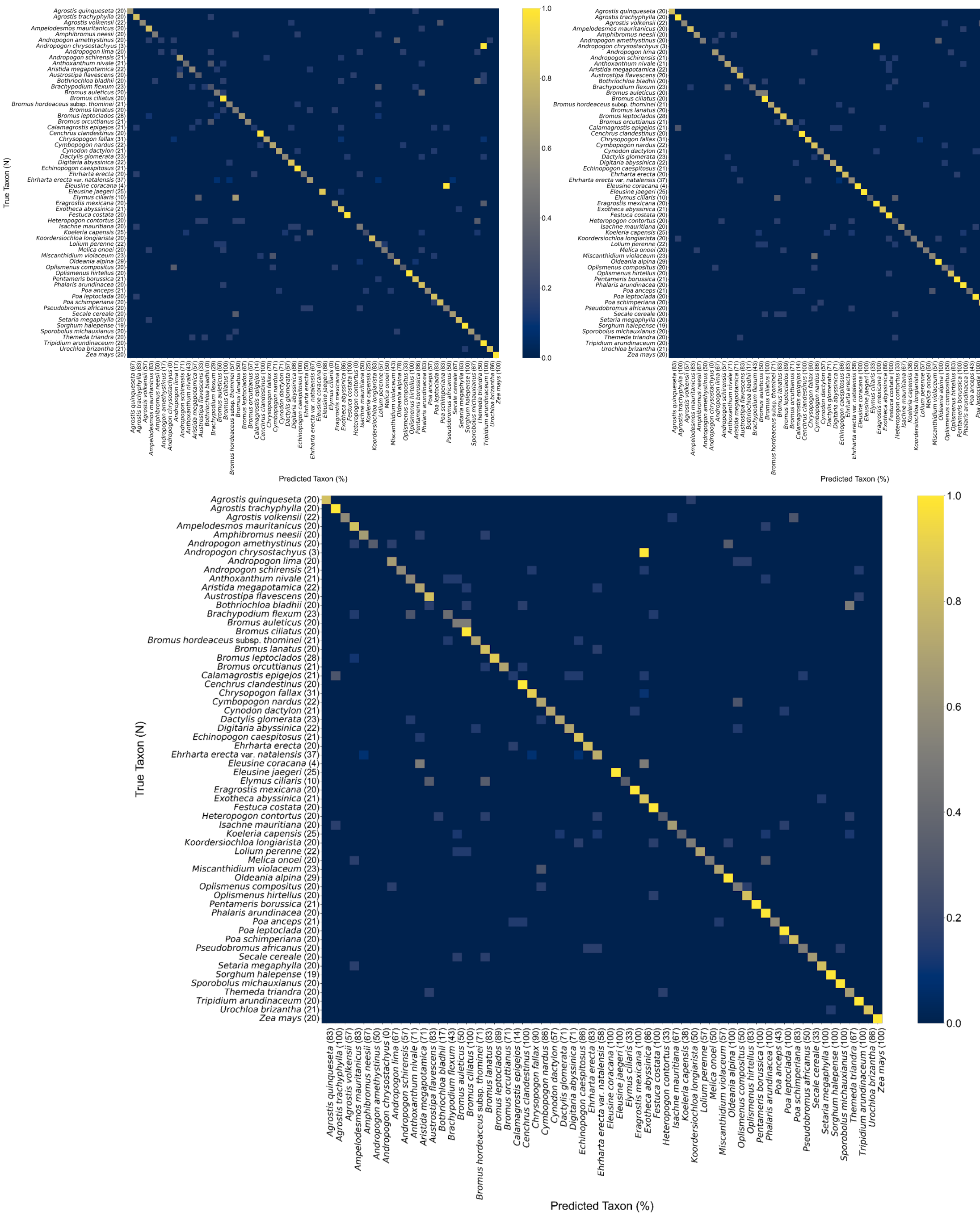

Agrostis quinqueseta (20)
Agrostis trachyphylla (20)
Agrostis volkensii (22)
Ampelodesmos mauritanicus (20)
Amphibromus neesii (20)
Andropogon amethystinus (20)
Andropogon chrysostachyus (3)
Andropogon lima (20)
Andropogon schirensis (21)
Anthoxanthum nivale (21)
Aristida megapotamica (22)
Austrostipa flavescens (20)
Bothriochloa bladhii (20)
Brachypodium flexum (23)
Bromus auleticus (20)
Bromus ciliatus (20)
Bromus hordeaceus subsp. thominei (21)
Bromus lanatus (20)
Bromus leptoclados (28)
Bromus orcuttianus (21)
Calamagrostis epigejos (21)
Cenchrus clandestinus (20)
Chrysopogon fallax (31)
Cymbopogon nardus (22)
Cynodon dactylon (21)
Dactylis glomerata (23)
Digitaria abyssinica (22)
Echinopogon caespitosus (21)
Ehrharta erecta (20)
Ehrharta erecta var. natalensis (37)
Eleusine coracana (4)
Eleusine jaegeri (25)
Elymus ciliaris (10)
Eragrostis mexicana (20)
Exotheca abyssinica (21)
Festuca costata (20)
Heteropogon contortus (20)
Isachne mauritiana (20)
Koeleria capensis (25)
Koordersiochloa longiarista (20)
Lolium perenne (22)
Melica onoei (20)
Miscanthidium violaceum (23)
Oldeania alpina (29)
Oplismenus compositus (20)
Oplismenus hirtellus (20)
Pentameris borussica (21)
Phalaris arundinacea (20)
Poa anceps (21)
Poa leptoclada (20)
Poa schimperiana (20)
Pseudobromus africanus (20)
Secale cereale (20)
Setaria megaphylla (20)
Sorghum halepense (19)
Sporobolus michauxianus (20)
Themeda triandra (20)
Tripidium arundinaceum (20)
Urochloa brizantha (21)
Zea mays (20)
True Taxon (N)
Agrostis quinqueseta (83)
Agrostis trachyphylla (100)
Agrostis volkensii (57)
Ampelodesmos mauritanicus (83)
Amphibromus neesii (67)
Andropogon amethystinus (50)
Andropogon chrysostachyus (0)
Andropogon lima (67)
Andropogon schirensis (57)
Anthoxanthum nivale (71)
Aristida megapotamica (71)
Austrostipa flavescens (83)
Bothriochloa bladhii (17)
Brachypodium flexum (43)
Bromus auleticus (50)
Bromus ciliatus (100)
Bromus hordeaceus subsp. thominei (71)
Bromus lanatus (83)
Bromus leptoclados (89)
Bromus orcuttianus (71)
Calamagrostis epigejos (14)
Cenchrus clandestinus (100)
Chrysopogon fallax (90)
Cymbopogon nardus (86)
Cynodon dactylon (57)
Dactylis glomerata (71)
Digitaria abyssinica (71)
Echinopogon caespitosus (86)
Ehrharta erecta (83)
Ehrharta erecta var. natalensis (58)
Eleusine coracana (100)
Eleusine jaegeri (100)
Elymus ciliaris (33)
Eragrostis mexicana (100)
Exotheca abyssinica (86)
Festuca costata (100)
Heteropogon contortus (33)
Isachne mauritiana (67)
Koeleria capensis (38)
Koordersiochloa longiarista (50)
Lolium perenne (57)
Melica onoei (50)
Miscanthidium violaceum (57)
Oldeania alpina (100)
Oplismenus compositus (50)
Oplismenus hirtellus (83)
Pentameris borussica (100)
Phalaris arundinacea (100)
Poa anceps (43)
Poa leptoclada (100)
Poa schimperiana (83)
Pseudobromus africanus (50)
Secale cereale (33)
Setaria megaphylla (100)
Sorghum halepense (100)
Sporobolus michauxianus (100)
Themeda triandra (67)
Tripidium arundinaceum (100)
Urochloa brizantha (86)
Zea mays (100)
Predicted Taxon (%)
0.0
0.2
0.4
0.6
0.8
1.0

### Average across five splits

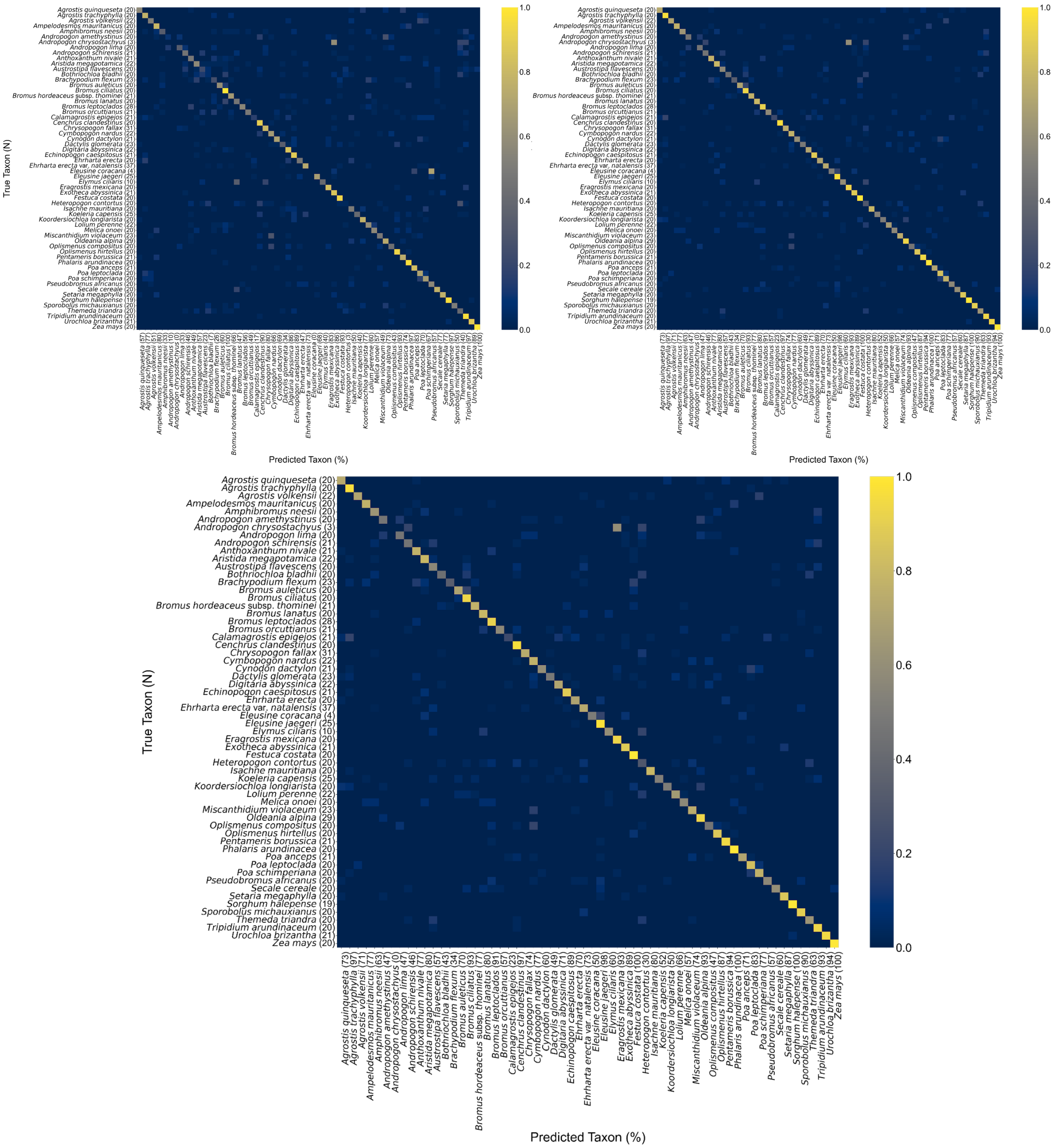

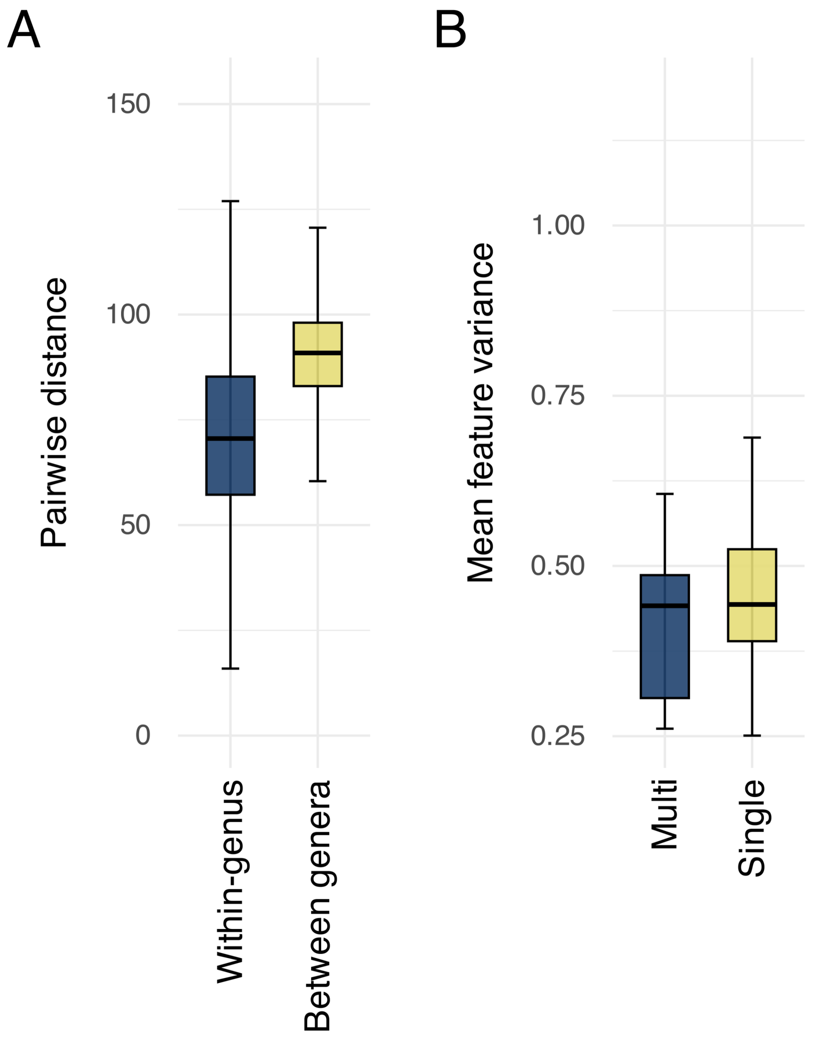


Fig. S2. Variability in pollen morphology captured by convolutional neural network (CNN) features. (A) Boxplots comparing pairwise Euclidean distances between standardized specimen embeddings within genera versus between genera. The results indicate that within-genus distances are smaller, showing genus-level clustering of pollen morphology. (B) Boxplots comparing within-species dispersion (mean feature variance per species) for taxa represented by multiple individuals ("Multi") versus a single individual ("Single"). The results indicate no significant difference between the two groups. All embeddings were derived from CNNs trained on superresolution images of modern Poaceae pollen. Boxplots show medians (lines), interquartile ranges (IQRs; boxes), and whiskers extending to the most extreme data points within 1.5×IQR from the hinges.

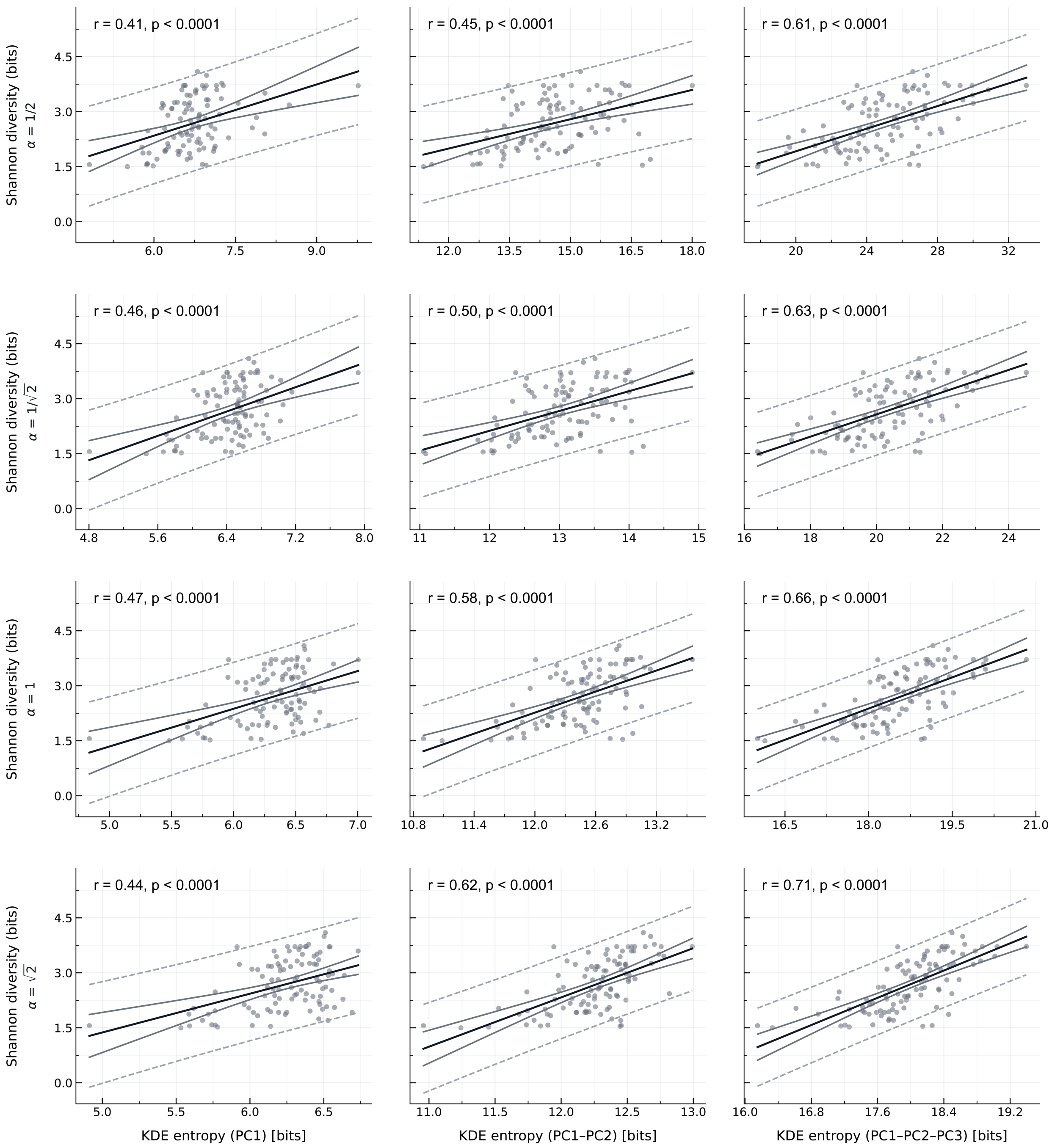


Fig. S3. Scatterplots depicting the correlation between Shannon diversity (y-axis) and Shannon entropy (x-axis) across 100 simulated grass pollen communities. Each community consisted of 50 grains drawn from the modern test dataset. Shannon diversity (in bits) was computed directly from species counts. KDE-entropy (in bits) was estimated from kernel density estimates applied to principal component analysis (PCA) scores of specimen features, using leave-one-out cross-validated bandwidths. Results are shown separately for the first principal component (PC), the first two PCs, and the first three PCs. Rows correspond to scaled bandwidths $\alpha h^*$, with scaling factors $\alpha \in \left\{\frac{1}{2}, \frac{1}{\sqrt{2}}, 1, \sqrt{2}\right\}$, illustrating robustness to smoothness. Ordinary least squares regression lines show the direction and strength of each relationship. Solid and dashed lines indicate 95% confidence and prediction intervals, respectively. In each panel, we report Pearson's correlation coefficient (r) and corresponding p-value.

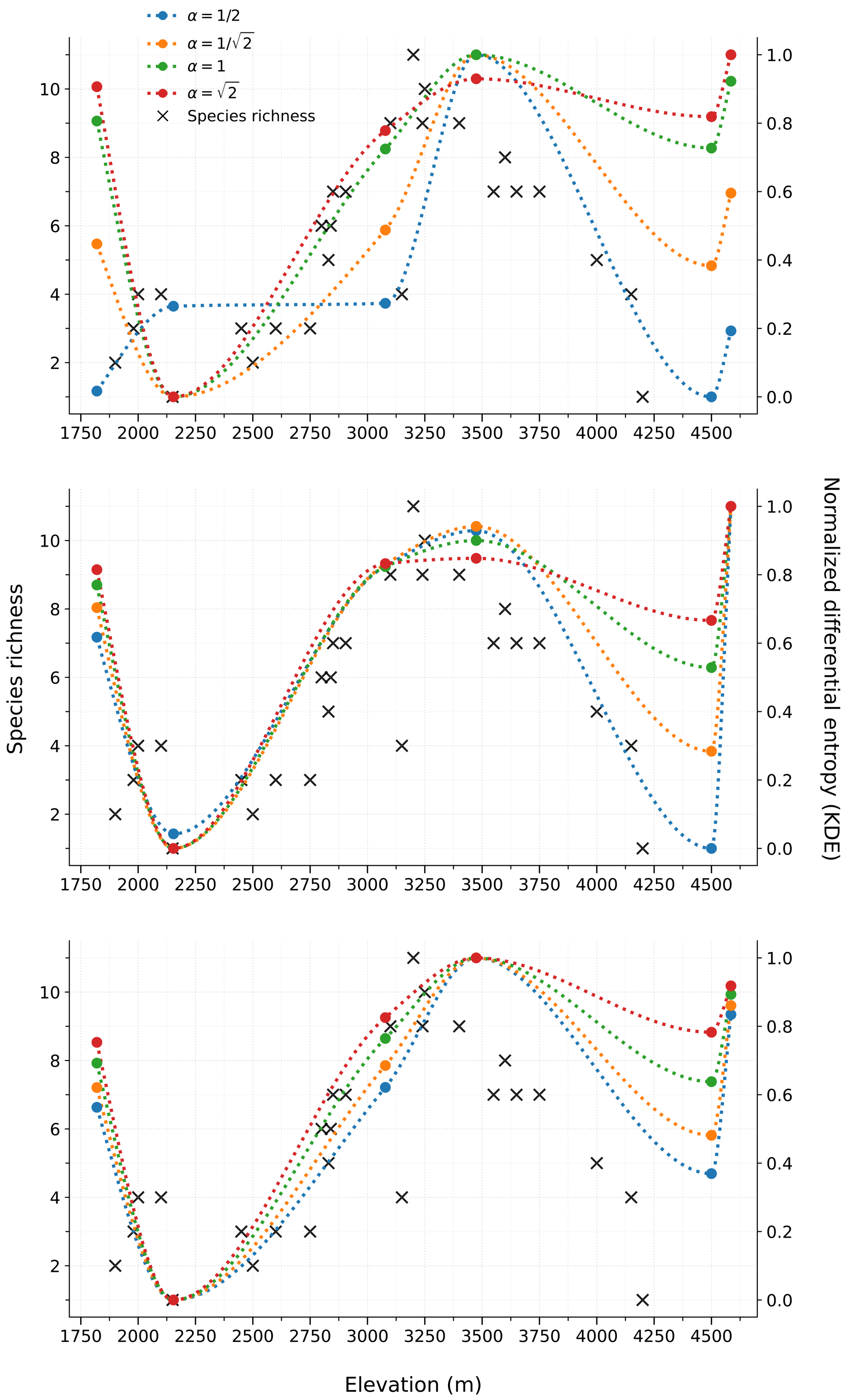


Fig. S4. Species richness from an elevational vegetation survey along Mt. Kenya (11), compared with normalized differential Shannon entropy estimates (used here as a proxy for diversity) derived from surface-sediment pollen assemblages from six lakes spanning the same elevational gradient. Gray points indicate the reference species richness at each elevation. Colored points and dotted lines indicate Shannon entropy estimates, computed as differential entropy of kernel density estimates (KDE) fitted to specimen scores in PC1, PC1-PC2, and PC1-PC2-PC3, respectively. Smoothing was applied using a piecewise cubic Hermite interpolating polynomial (PCHIP) (21) to produce continuous, non-negative curves that preserve the local shape of the data.

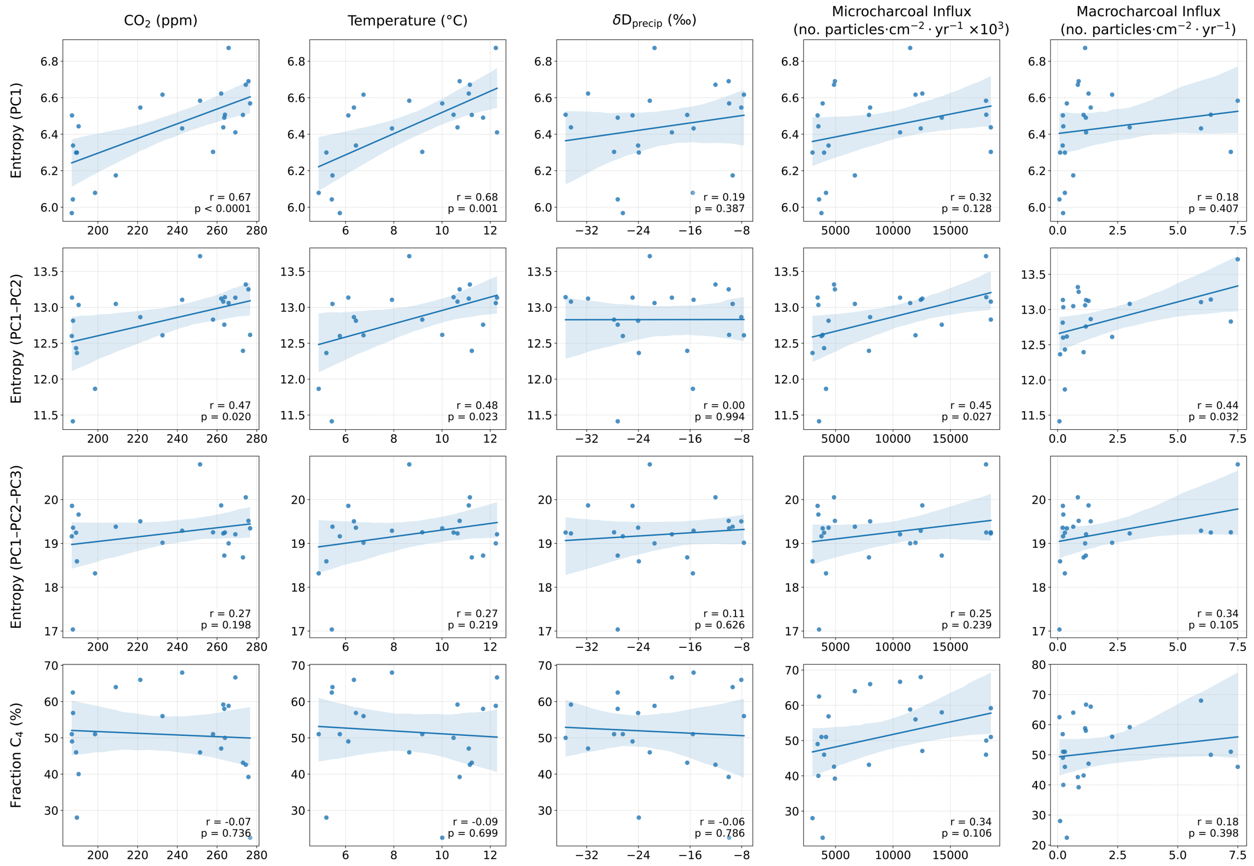


Fig. S5. Correlation of Shannon entropy (along PC1, PC1-PC2, and PC1-PC2-PC3) and inferred proportion of $C_4$ grasses of the Lake Rutundu record with five paleoenvironmental variables. From left to right, atmospheric $CO_2$ concentration, reconstructed local temperature, reconstructed local precipitation ($\delta D_{precip}$), and micro- and macrocharcoal fluxes as proxies of fire prevalence. Linear regression lines are fitted to each plot, and shaded areas indicate the 95% confidence intervals. Pearson correlation coefficients (r) and associated p-values are shown within each panel.

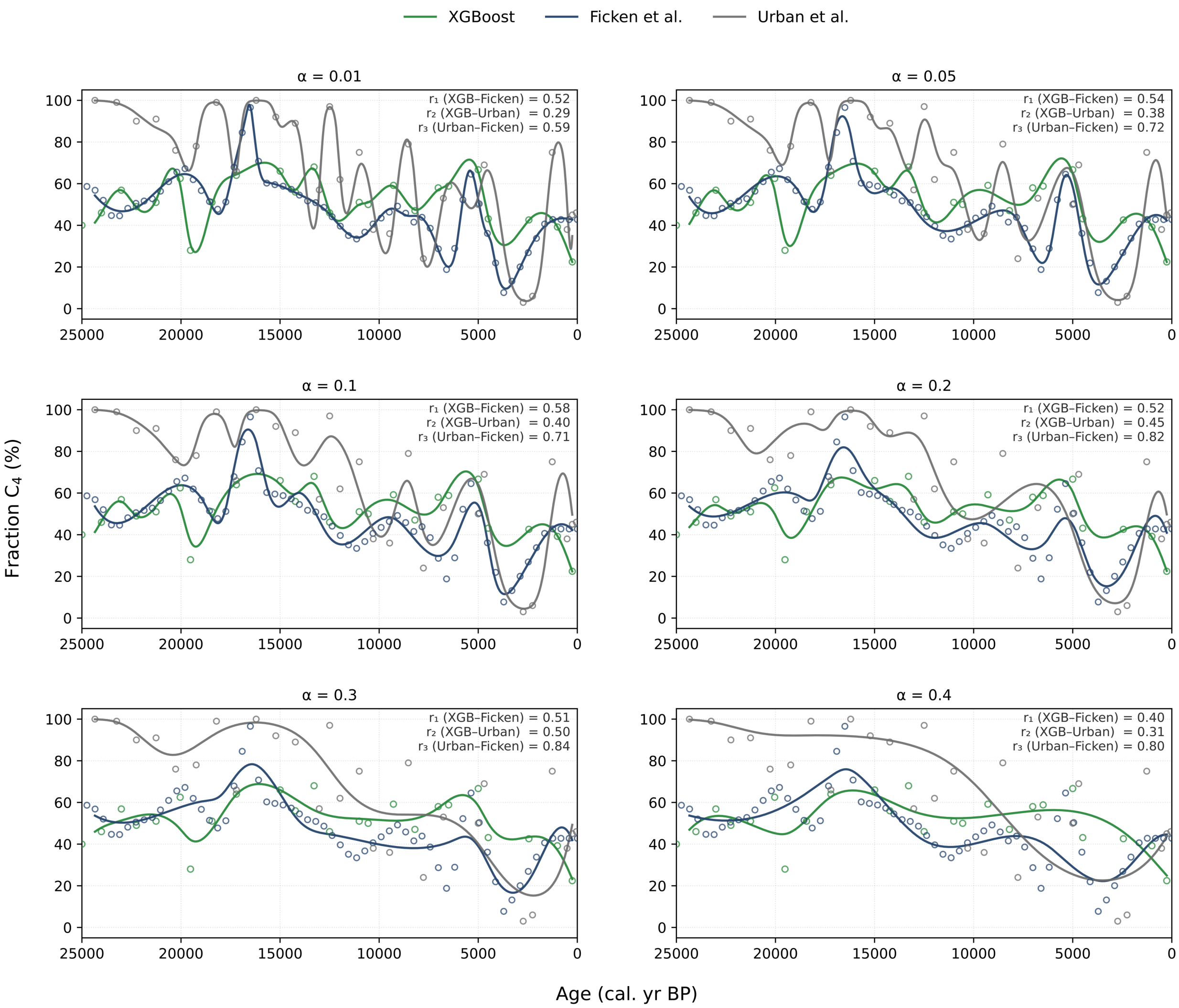


Fig. S6. Comparison of $C_4$ reconstructions from this study (XGBoost classifier, green), Ficken et al. (2002) (blue), and Urban et al. (2015) (gray) smoothed with cubic splines using different smoothing factors (from $\alpha$ = 0.01 to $\alpha$ = 0.4). Open circles denote the underlying series: green circles are individual XGBoost predictions; blue circles are values used to construct the midpoint series derived from the published upper and lower bounds of Ficken et al. (2002); gray circles are $\delta^{13}C$-based estimates from Urban et al. (2015). The three reconstructions show broadly similar temporal patterns, with a general decline in $C_4$ grass abundance from ~25 Ka BP to the present. Our estimates exhibit fair correlation with both independent proxies and closely match the absolute values and long-term trajectories of Ficken et al.'s midpoint series.

Table S1. Modern grass species sampled for pollen from herbarium specimens, including number of pollen grains analyzed and photosynthetic pathway classification. Species that included multiple individuals are presented with an asterisk (*).

| **Species** | **Number of pollen grains** | **Photosynthetic Pathway** | **Collection** |
|---|---|---|---|
| *Agrostis quinqueseta* | 20 | $C_3$ | MOBOT |
| *Agrostis trachyphylla* | 20 | $C_3$ | MOBOT |
| *Agrostis volkensii* | 22 | $C_3$ | MOBOT |
| *Ampelodesmos mauritanicus** | 20 | $C_3$ | STRI |
| *Amphibromus neesii* | 20 | $C_3$ | Harvard |
| *Andropogon amethystinus* | 20 | $C_4$ | MOBOT |
| *Andropogon chrysostachyus* | 3 | $C_4$ | Harvard |
| *Andropogon lima* | 20 | $C_4$ | MOBOT |
| *Andropogon schirensis* | 21 | $C_4$ | MOBOT |
| *Anthoxanthum nivale* | 21 | $C_3$ | MOBOT |
| *Aristida megapotamica* | 22 | $C_4$ | Harvard |
| *Austrostipa flavescens* | 20 | $C_3$ | Harvard |
| *Bothriochloa bladhii* | 20 | $C_4$ | Harvard |
| *Brachypodium flexum* | 23 | $C_3$ | MOBOT |
| *Bromus auleticus* | 20 | $C_3$ | Harvard |
| *Bromus ciliatus* | 20 | $C_3$ | Harvard |
| *Bromus hordeaceus* subsp. *thominei* | 21 | $C_3$ | Harvard |
| *Bromus lanatus* | 20 | $C_3$ | Harvard |
| *Bromus leptoclados* | 28 | $C_3$ | MOBOT |
| *Bromus orcuttianus* | 21 | $C_3$ | Harvard |
| *Calamagrostis epigejos* | 21 | $C_3$ | MOBOT |
| *Cenchrus clandestinus* | 20 | $C_4$ | MOBOT |

| | | | |
|---|---|---|---|
| *Chrysopogon fallax* | 31 | $C_4$ | Harvard |
| *Cymbopogon nardus* | 22 | $C_4$ | MOBOT |
| *Cynodon dactylon* | 21 | $C_4$ | Harvard |
| *Dactylis glomerata** | 23 | $C_3$ | Sigma |
| *Digitaria abyssinica* | 22 | $C_4$ | MOBOT |
| *Echinopogon caespitosus* | 21 | $C_3$ | Harvard |
| *Ehrharta erecta* | 20 | $C_3$ | MOBOT |
| *Ehrharta erecta* var. *natalensis* | 37 | $C_3$ | Harvard |
| *Eleusine coracana* | 4 | $C_4$ | Harvard |
| *Eleusine jaegeri* | 25 | $C_4$ | MOBOT |
| *Elymus ciliaris* | 10 | $C_3$ | Harvard |
| *Eragrostis mexicana* | 20 | $C_4$ | Harvard |
| *Exotheca abyssinica* | 21 | $C_4$ | MOBOT |
| *Festuca costata* | 20 | $C_3$ | MOBOT |
| *Heteropogon contortus* | 20 | $C_4$ | Harvard |
| *Isachne mauritiana* | 20 | $C_3$ | MOBOT |
| *Koeleria capensis* | 25 | $C_3$ | MOBOT |
| *Koordersiochloa longiarista* | 20 | $C_3$ | MOBOT |
| *Lolium perenne* | 22 | $C_3$ | Harvard |
| *Melica onoei* | 20 | $C_3$ | Harvard |
| *Miscanthidium violaceum* | 23 | $C_4$ | MOBOT |
| *Oldeania alpina* | 29 | $C_3$ | MOBOT |
| *Oplismenus compositus* | 20 | $C_3$ | Harvard |
| *Oplismenus hirtellus* | 20 | $C_3$ | Harvard |
| *Pentameris borussica* | 21 | $C_3$ | MOBOT |
| *Phalaris arundinacea* | 20 | $C_3$ | MOBOT |
| *Poa anceps* | 21 | $C_3$ | Harvard |

| | | | |
|---|---|---|---|
| *Poa leptoclada* | 20 | $C_3$ | Harvard |
| *Poa schimperiana* | 20 | $C_3$ | MOBOT |
| *Pseudobromus africanus* | 20 | $C_3$ | MOBOT |
| *Secale cereale** | 20 | $C_3$ | Sigma |
| *Setaria megaphylla* | 20 | $C_4$ | MOBOT |
| *Sorghum halepense** | 19 | $C_4$ | Sigma |
| *Sporobolus michauxianus** | 20 | $C_4$ | STRI |
| *Themeda triandra* | 20 | $C_4$ | Harvard |
| *Tripidium arundinaceum* | 20 | $C_4$ | Harvard |
| *Urochloa brizantha* | 21 | $C_4$ | Harvard |
| *Zea mays** | 20 | $C_4$ | STRI |

Table S2. Native range of the 60 grass species represented in our modern grass pollen dataset, based on (22). Species present on Mt. Kenya, as documented in (23), are marked with an asterisk (*), while species not recorded on Mt. Kenya but belonging to genera that are present there are marked with a dagger (†).

| Species | Native Range |
|---|---|
| *Agrostis quinqueseta** | Cameroon, Ethiopia, Kenya, Rwanda, Tanzania, Uganda, Zaïre |
| *Agrostis trachyphylla** | Kenya, Tanzania, Uganda, Zaïre |
| *Agrostis volkensii** | Ethiopia, Kenya, Tanzania, Uganda |
| *Ampelodesmos mauritanicus* | Algeria, Baleares, Corse, France, Greece, Italy, Libya, Morocco, Sardegna, Sicilia, Spain, Tunisia |
| *Amphibromus neesii* | New South Wales, Tasmania, Victoria |
| *Andropogon amethystinus** | Cameroon, Cape Provinces, Eritrea, Ethiopia, Free State, Gulf of Guinea Is., India, Kenya, KwaZulu-Natal, Lesotho, Malawi, Myanmar, Nigeria, Rwanda, Somalia, Sudan, Tanzania, Togo, Uganda, Yemen, Zambia, Zaïre |
| *Andropogon chrysostachyus** | Ethiopia, Kenya, Tanzania |
| *Andropogon lima** | Cameroon, Ethiopia, Kenya, Malawi, Rwanda, Sudan, Tanzania, Uganda |
| *Andropogon schirensis** | Angola, Benin, Botswana, Burkina, Burundi, Cameroon, Cape Provinces, Central African Republic, Chad, Congo, Ethiopia, Free State, Gabon, Ghana, Guinea, Ivory Coast, Kenya, KwaZulu-Natal, Lesotho, Malawi, Mali, Mozambique, Namibia, Niger, Nigeria, Northern Provinces, Rwanda, Senegal, Sierra Leone, Sudan, Swaziland, Tanzania, Togo, Uganda, Zambia, Zaïre, Zimbabwe |
| *Anthoxanthum nivale** | Kenya, Rwanda, Tanzania, Uganda, Zaïre |
| *Aristida megapotamica*† | Argentina Northeast, Belize, Bolivia, Brazil Northeast, Brazil South, Brazil Southeast, Brazil West-Central, Colombia, El Salvador, Honduras, Paraguay, Peru, Uruguay |
| *Austrostipa flavescens* | New South Wales, South Australia, Tasmania, Victoria, Western Australia |

| | |
|---|---|
| *Bothriochloa bladhii*† | Afghanistan, Angola, Bangladesh, Benin, Borneo, Botswana, Burkina, Burundi, Cameroon, Cape Provinces, Cape Verde, Chad, China North-Central, China South-Central, China Southeast, East Himalaya, Ethiopia, Ghana, Hainan, India, Iran, Ivory Coast, Jawa, Kazakhstan, Kenya, Kirgizstan, KwaZulu-Natal, Laos, Lesser Sunda Is., Madagascar, Malawi, Malaya, Mali, Maluku, Mauritius, Mozambique, Myanmar, Namibia, Nansei-shoto, Nepal, New Guinea, New South Wales, Nigeria, North Caucasus, Northern Provinces, Northern Territory, Oman, Pakistan, Philippines, Queensland, Rodrigues, Réunion, Senegal, South Australia, South China Sea, Sri Lanka, Sudan, Sulawesi, Sumatera, Swaziland, Tadzhikistan, Taiwan, Tanzania, Thailand, Transcaucasus, Turkey, Uganda, Uzbekistan, Vietnam, Wallis-Futuna Is., West Himalaya, Western Australia, Xinjiang, Yemen, Zambia, Zaïre, Zimbabwe |
| *Brachypodium flexum** | Burundi, Cabinda, Cameroon, Cape Provinces, Eritrea, Ethiopia, Free State, Gulf of Guinea Is., Kenya, KwaZulu-Natal, Lesotho, Madagascar, Malawi, Mozambique, Nigeria, Northern Provinces, Rwanda, Sierra Leone, Sudan, Swaziland, Tanzania, Uganda, Zambia, Zimbabwe |
| *Bromus auleticus*† | Argentina Northeast, Argentina Northwest, Brazil South, Uruguay |
| *Bromus ciliatus*† | Alabama, Alaska, Alberta, Aleutian Is., Arizona, British Columbia, California, Colorado, Connecticut, Idaho, Illinois, Indiana, Inner Mongolia, Iowa, Japan, Kamchatka, Khabarovsk, Korea, Kuril Is., Labrador, Magadan, Maine, Manitoba, Maryland, Massachusetts, Minnesota, Mongolia, Montana, Nebraska, New Brunswick, New Hampshire, New Mexico, New York, Newfoundland, North Carolina, North Dakota, Northwest Territories, Nova Scotia, Ohio, Ontario, Oregon, Pennsylvania, Prince Edward I., Québec, Rhode I., Sakhalin, Saskatchewan, South Dakota, Tennessee, Texas, Utah, Vermont, Virginia, Washington, West Virginia, Wisconsin, Wyoming, Yukon |
| *Bromus hordeaceus* subsp. *thominei*† | Austria, Azores, Baleares, Belgium, Corse, Denmark, France, Germany, Great Britain, Greece, Ireland, Italy, Morocco, Netherlands, Norway, Portugal, Sardegna, Spain, Sweden, Turkey |

| | |
|---|---|
| *Bromus lanatus*[†] | Argentina Northwest, Bolivia, Chile North, Colombia, Ecuador, Peru, Venezuela |
| *Bromus leptoclados** | Burundi, Cameroon, Cape Provinces, Eritrea, Ethiopia, Free State, Gulf of Guinea Is., Kenya, KwaZulu-Natal, Lesotho, Malawi, Northern Provinces, Rwanda, Réunion, Somalia, Sudan, Tanzania, Uganda, Yemen, Zimbabwe |
| *Bromus orcuttianus*[†] | Arizona, California, Nevada, Oregon, Washington |
| *Calamagrostis epigejos** | Afghanistan, Albania, Altay, Amur, Austria, Baltic States, Belarus, Belgium, Bulgaria, Buryatiya, Cape Provinces, Central European Russia, China North-Central, Chita, Corse, Cyprus, Czechoslovakia, Denmark, East European Russia, Ethiopia, Finland, France, Germany, Great Britain, Greece, Hungary, Inner Mongolia, Iran, Iraq, Ireland, Irkutsk, Italy, Japan, Kazakhstan, Kenya, Khabarovsk, Kirgizstan, Korea, Krasnoyarsk, Krym, Kuril Is., Lebanon-Syria, Manchuria, Mongolia, Nepal, Netherlands, North Caucasus, North European Russia, Northern Provinces, Northwest European Russia, Norway, Pakistan, Poland, Primorye, Romania, Rwanda, Sakhalin, Sardegna, Sicilia, South European Russia, Spain, Sudan, Sweden, Switzerland, Tadzhikistan, Tanzania, Transcaucasus, Turkey, Turkey-in-Europe, Turkmenistan, Tuva, Uganda, Ukraine, Uzbekistan, West Himalaya, West Siberia, Xinjiang, Yakutskiya, Yugoslavia |
| *Cenchrus clandestinus** | Burundi, Eritrea, Ethiopia, Kenya, Malawi, Rwanda, Tanzania, Uganda, Zaïre, Zimbabwe |
| *Chrysopogon fallax* | New South Wales, Northern Territory, Queensland, South Australia, Victoria, Western Australia |
| *Cymbopogon nardus** | Angola, Assam, Bangladesh, Botswana, Burundi, Cambodia, Cape Provinces, East Himalaya, Free State, India, Kenya, KwaZulu-Natal, Laos, Lesotho, Madagascar, Mozambique, Myanmar, Northern Provinces, Rwanda, Seychelles, Sri Lanka, Sudan, Swaziland, Tanzania, Uganda, Vietnam, West Himalaya, Zaïre, Zimbabwe |
| *Cynodon dactylon** | Afghanistan, Albania, Aldabra, Algeria, Andaman Is., Angola, Assam, Austria, Azores, Baleares, Bangladesh, Benin, Bismarck Archipelago, Borneo, Botswana, Bulgaria, Burkina, Burundi, Cambodia, Cameroon, Canary Is., Cape Provinces, Cape Verde, Caprivi Strip, Central African Republic, Central |

| | |
|---|---|
| | European Russia, Chad, China North-Central, China South-Central, China Southeast, Congo, Corse, Cyprus, Czechoslovakia, Djibouti, East Aegean Is., East Himalaya, Egypt, Equatorial Guinea, Eritrea, Ethiopia, France, Free State, Gabon, Gambia, Ghana, Great Britain, Greece, Guinea, Guinea-Bissau, Gulf of Guinea Is., Gulf States, Hainan, Hungary, India, Iran, Iraq, Italy, Ivory Coast, Japan, Jawa, Kazakhstan, Kenya, Kirgizstan, Korea, Kriti, Krym, Kuwait, KwaZulu-Natal, Laccadive Is., Laos, Lebanon-Syria, Lesotho, Lesser Sunda Is., Liberia, Libya, Madagascar, Madeira, Malawi, Malaya, Mali, Maluku, Mauritania, Mauritius, Morocco, Mozambique, Mozambique Channel Is., Myanmar, Namibia, Nansei-shoto, Nepal, New Guinea, New South Wales, Nicobar Is., Niger, Nigeria, North Caucasus, Northern Provinces, Northern Territory, Ogasawara-shoto, Oman, Pakistan, Palestine, Philippines, Portugal, Queensland, Rodrigues, Romania, Rwanda, Réunion, Sardegna, Saudi Arabia, Senegal, Seychelles, Sicilia, Sierra Leone, Sinai, Socotra, Solomon Is., Somalia, South Australia, South China Sea, South European Russia, Spain, Sri Lanka, Sudan, Sumatera, Swaziland, Switzerland, Tadzhikistan, Taiwan, Tanzania, Tasmania, Thailand, Togo, Transcaucasus, Tunisia, Turkey, Turkey-in-Europe, Turkmenistan, Uganda, Ukraine, Uzbekistan, Victoria, Vietnam, West Himalaya, Western Australia, Western Sahara, Yemen, Yugoslavia, Zambia, Zaïre, Zimbabwe |
| *Dactylis glomerata** | Afghanistan, Albania, Algeria, Altay, Assam, Austria, Azores, Baleares, Baltic States, Belarus, Belgium, Bulgaria, Buryatiya, Canary Is., Central European Russia, China North-Central, China South-Central, China Southeast, Corse, Cyprus, Czechoslovakia, Denmark, East Aegean Is., East European Russia, East Himalaya, Egypt, Finland, France, Føroyar, Germany, Great Britain, Greece, Hungary, Iceland, India, Inner Mongolia, Iran, Iraq, Ireland, Irkutsk, Italy, Kazakhstan, Kirgizstan, Krasnoyarsk, Kriti, Krym, Lebanon-Syria, Libya, Madeira, Mongolia, Morocco, Nepal, Netherlands, North Caucasus, North European Russia, Northwest European Russia, Norway, Pakistan, Palestine, Poland, Portugal, Romania, Sardegna, Sicilia, South European Russia, Spain, Sri Lanka, Sweden, Switzerland, Tadzhikistan, Taiwan, Tibet, Transcaucasus, Tunisia, Turkey, Turkey-in-Europe, Turkmenistan, Tuva, Ukraine, Uzbekistan, West Himalaya, West Siberia, Xinjiang, |

| | |
|---|---|
| | Yugoslavia |
| *Digitaria abyssinica** | Botswana, Burundi, Cameroon, Cape Provinces, Comoros, Congo, Djibouti, Eritrea, Ethiopia, Free State, Gabon, Kenya, KwaZulu-Natal, Lesotho, Malawi, Malaya, Mozambique, Namibia, New Guinea, Nigeria, Northern Provinces, Rwanda, Réunion, Saudi Arabia, Seychelles, Somalia, Sri Lanka, Sudan, Swaziland, Tanzania, Uganda, Vietnam, Yemen, Zambia, Zaïre, Zimbabwe |
| *Echinopogon caespitosus* | New South Wales, Queensland, Victoria |
| *Ehrharta erecta** | Botswana, Cape Provinces, Eritrea, Ethiopia, Free State, Kenya, KwaZulu-Natal, Lesotho, Malawi, Mozambique, Northern Provinces, Rwanda, Réunion, Saudi Arabia, Somalia, Sudan, Swaziland, Tanzania, Uganda, Yemen, Zambia, Zaïre, Zimbabwe |
| *Ehrharta erecta* var. *natalensis** | Botswana, Cape Provinces, Eritrea, Ethiopia, Free State, Kenya, KwaZulu-Natal, Lesotho, Malawi, Mozambique, Northern Provinces, Rwanda, Réunion, Saudi Arabia, Somalia, Sudan, Swaziland, Tanzania, Uganda, Yemen, Zambia, Zaïre, Zimbabwe |
| *Eleusine coracana*† | Angola, Botswana, Burkina, Burundi, Cameroon, Cape Provinces, Chad, Comoros, Egypt, Eritrea, Ethiopia, Free State, Gambia, Ghana, Guinea-Bissau, Kenya, KwaZulu-Natal, Lesotho, Madagascar, Malawi, Mali, Mozambique, Namibia, Niger, Nigeria, Northern Provinces, Oman, Rwanda, Saudi Arabia, Senegal, Sierra Leone, Sinai, Socotra, Somalia, Sudan, Swaziland, Tanzania, Togo, Uganda, Yemen, Zambia, Zaïre, Zimbabwe |
| *Eleusine jaegeri** | Ethiopia, Kenya, Tanzania, Uganda |
| *Elymus ciliaris*† | Assam, China North-Central, China South-Central, China Southeast, Inner Mongolia, Japan, Khabarovsk, Korea, Manchuria, Mongolia, Nansei-shoto, Primorye, Taiwan |
| *Eragrostis mexicana*† | Argentina Northeast, Argentina Northwest, Argentina South, Arizona, Bolivia, Brazil North, Brazil Northeast, Brazil South, Brazil Southeast, Brazil West-Central, British Columbia, California, Chile North, Colombia, Colorado, Costa Rica, Ecuador, El Salvador, Galápagos, Guatemala, Honduras, Mexico Central, Mexico Gulf, Mexico Northeast, Mexico Northwest, Mexico Southeast, Mexico Southwest, Nevada, New Mexico, Nicaragua, Oklahoma, |

| | |
|---|---|
| | Oregon, Panamá, Peru, Texas, Uruguay, Utah, Venezuela, Washington |
| *Exotheca abyssinica** | Burundi, Eritrea, Ethiopia, Kenya, Malawi, Mozambique, Rwanda, Sudan, Tanzania, Uganda, Vietnam, Zambia, Zaïre |
| *Festuca costata** | Angola, Cape Provinces, Free State, Kenya, KwaZulu-Natal, Lesotho, Malawi, Mozambique, Northern Provinces, Swaziland, Tanzania, Zimbabwe |
| *Heteropogon contortus* | Afghanistan, Albania, Algeria, Andaman Is., Angola, Argentina Northeast, Argentina Northwest, Arizona, Assam, Baleares, Bangladesh, Belize, Benin, Bolivia, Botswana, Brazil North, Burkina, Burundi, California, Cambodia, Cameroon, Canary Is., Cape Provinces, Cape Verde, Caprivi Strip, Central African Republic, Chad, China North-Central, China South-Central, China Southeast, Colombia, Comoros, Cuba, Djibouti, Dominican Republic, East Himalaya, Ecuador, El Salvador, Eritrea, Ethiopia, Fiji, Florida, France, Free State, Ghana, Guatemala, Gulf of Guinea Is., Guyana, Hainan, Haiti, Hawaii, Honduras, India, Iran, Iraq, Italy, Ivory Coast, Jamaica, Japan, Jawa, Kenya, KwaZulu-Natal, Laccadive Is., Lebanon-Syria, Leeward Is., Lesotho, Lesser Sunda Is., Madagascar, Malawi, Malaya, Mali, Maluku, Marianas, Marquesas, Mauritania, Mauritius, Mexican Pacific Is., Mexico Central, Mexico Gulf, Mexico Northeast, Mexico Northwest, Mexico Southeast, Mexico Southwest, Morocco, Mozambique, Myanmar, Namibia, Nansei-shoto, Nepal, Netherlands Antilles, New Caledonia, New Guinea, New Mexico, Nicaragua, Nicobar Is., Niger, Nigeria, Northern Provinces, Oman, Pakistan, Paraguay, Peru, Philippines, Puerto Rico, Rodrigues, Rwanda, Réunion, Samoa, Saudi Arabia, Senegal, Seychelles, Sicilia, Sierra Leone, Society Is., Socotra, Somalia, South China Sea, Spain, Sri Lanka, Sudan, Sulawesi, Sumatera, Swaziland, Switzerland, Taiwan, Tanzania, Texas, Thailand, Tibet, Togo, Tonga, Tunisia, Uganda, Vietnam, West Himalaya, Windward Is., Yemen, Yugoslavia, Zambia, Zaïre, Zimbabwe |
| *Isachne mauritiana** | Burundi, Cameroon, Congo, Ghana, Gulf of Guinea Is., Kenya, Madagascar, Malawi, Mauritius, Mozambique, Nigeria, Rwanda, Réunion, Sudan, Tanzania, Uganda, Zambia, Zaïre, Zimbabwe |

| | |
|---|---|
| *Koeleria capensis** | Cameroon, Cape Provinces, Ethiopia, Free State, Kenya, KwaZulu-Natal, Lesotho, Malawi, Mozambique, Northern Provinces, Sudan, Swaziland, Tanzania, Uganda, Yemen, Zambia, Zimbabwe |
| *Koordersiochloa longiarista** | Cameroon, Cape Provinces, Ethiopia, Gulf of Guinea Is., Jawa, Kenya, KwaZulu-Natal, Lesser Sunda Is., Malawi, Nigeria, Philippines, Réunion, Sudan, Tanzania, Uganda, Zimbabwe |
| *Lolium perenne* | Afghanistan, Albania, Algeria, Austria, Azores, Baltic States, Belarus, Belgium, Bulgaria, Canary Is., Central European Russia, Corse, Cyprus, Czechoslovakia, Denmark, East Aegean Is., East European Russia, East Himalaya, Egypt, France, Germany, Great Britain, Greece, Hungary, Iran, Iraq, Ireland, Italy, Kazakhstan, Kirgizstan, Krasnoyarsk, Kriti, Krym, Lebanon-Syria, Libya, Madeira, Mauritania, Morocco, Nepal, Netherlands, North Caucasus, North European Russia, Northwest European Russia, Norway, Pakistan, Palestine, Poland, Portugal, Romania, Sardegna, Selvagens, Sicilia, South European Russia, Spain, Sweden, Switzerland, Tadzhikistan, Transcaucasus, Tunisia, Turkey, Turkey-in-Europe, Turkmenistan, Ukraine, West Himalaya, West Siberia, Yugoslavia |
| *Melica onoei* | China North-Central, China South-Central, China Southeast, East Himalaya, Inner Mongolia, Japan, Korea, Pakistan, Taiwan, Tibet, West Himalaya, Xinjiang |
| *Miscanthidium violaceum** | Burundi, Kenya, Rwanda, Tanzania, Uganda, Zambia, Zaïre |
| *Oldeania alpina** | Burundi, Cameroon, Congo, Ethiopia, Kenya, Malawi, Rwanda, Sudan, Tanzania, Uganda, Zambia, Zaïre |
| *Oplismenus compositus** | Alabama, Andaman Is., Argentina Northeast, Argentina Northwest, Arkansas, Assam, Bangladesh, Belize, Bermuda, Bismarck Archipelago, Borneo, Brazil South, Burundi, Cambodia, Caroline Is., China South-Central, China Southeast, Christmas I., Colombia, Comoros, Costa Rica, Cuba, Dominican Republic, East Himalaya, Ecuador, El Salvador, Equatorial Guinea, Eritrea, Ethiopia, Florida, Georgia, Guatemala, Hainan, Haiti, Honduras, India, Iran, Jamaica, Japan, Jawa, Kazan-retto, Kenya, Korea, Laccadive Is., Laos, Leeward Is., Lesser Sunda Is., Louisiana, Madagascar, Malawi, Malaya, Maldives, Maluku, Mauritius, Mexico Central, Mexico Gulf, |

| | |
|---|---|
| | Mexico Northeast, Mexico Northwest, Mexico Southeast, Mexico Southwest, Mississippi, Missouri, Mozambique, Myanmar, Nansei-shoto, Nepal, New Guinea, New South Wales, Nicaragua, Nicobar Is., North Carolina, Northern Territory, Ogasawara-shoto, Oklahoma, Oman, Pakistan, Panamá, Paraguay, Peru, Philippines, Puerto Rico, Queensland, Réunion, Seychelles, Socotra, South Carolina, Sri Lanka, Sudan, Sulawesi, Sumatera, Taiwan, Tanzania, Texas, Thailand, Tibet, Transcaucasus, Trinidad-Tobago, Uganda, Uruguay, Vanuatu, Vietnam, Virginia, West Himalaya, Windward Is., Yemen, Zambia, Zimbabwe |
| *Oplismenus hirtellus** | Angola, Argentina Northeast, Argentina Northwest, Bahamas, Belize, Benin, Bermuda, Bismarck Archipelago, Bolivia, Borneo, Botswana, Brazil North, Brazil South, Brazil Southeast, Brazil West-Central, Burkina, Burundi, Cameroon, Canary Is., Cape Provinces, Cape Verde, Caroline Is., Cayman Is., Central African Republic, China South-Central, China Southeast, Colombia, Comoros, Congo, Cook Is., Costa Rica, Cuba, Dominican Republic, Ecuador, El Salvador, Equatorial Guinea, Eritrea, Ethiopia, Fiji, Florida, French Guiana, Gabon, Galápagos, Gambia, Ghana, Guatemala, Guinea, Guinea-Bissau, Gulf of Guinea Is., Guyana, Haiti, Honduras, Ivory Coast, Jamaica, Japan, Jawa, Kenya, Kermadec Is., Korea, KwaZulu-Natal, Leeward Is., Lesser Sunda Is., Liberia, Madagascar, Madeira, Malawi, Mali, Maluku, Marianas, Marquesas, Mauritius, Mexican Pacific Is., Mexico Central, Mexico Gulf, Mexico Northeast, Mexico Northwest, Mexico Southeast, Mexico Southwest, Nansei-shoto, New Caledonia, New Guinea, New South Wales, New Zealand North, New Zealand South, Nicaragua, Nigeria, Niue, Norfolk Is., Northern Provinces, Northern Territory, Panamá, Paraguay, Peru, Philippines, Pitcairn Is., Puerto Rico, Queensland, Rodrigues, Rwanda, Réunion, Samoa, Senegal, Sierra Leone, Society Is., Solomon Is., Sudan, Sulawesi, Suriname, Swaziland, Taiwan, Tanzania, Thailand, Togo, Tonga, Trinidad-Tobago, Tuamotu, Tubuai Is., Uganda, Uruguay, Vanuatu, Venezuela, Venezuelan Antilles, Victoria, Vietnam, Wallis-Futuna Is., Western Australia, Windward Is., Yemen, Zambia, Zaïre, Zimbabwe |
| *Pentameris borussica** | Ethiopia, Kenya, Tanzania, Uganda |
| *Phalaris arundinacea** | Afghanistan, Alabama, Alaska, Albania, Alberta, |

| | |
|---|---|
| | Altay, Amur, Arizona, Arkansas, Austria, Baltic States, Belarus, Belgium, British Columbia, Bulgaria, Buryatiya, California, Central European Russia, China North-Central, China South-Central, China Southeast, Chita, Colorado, Connecticut, Corse, Czechoslovakia, Delaware, Denmark, District of Columbia, East European Russia, East Himalaya, Ethiopia, Finland, France, Føroyar, Germany, Great Britain, Greece, Hungary, Illinois, Indiana, Inner Mongolia, Iowa, Iran, Iraq, Ireland, Irkutsk, Italy, Japan, Kamchatka, Kansas, Kazakhstan, Kentucky, Kenya, Khabarovsk, Kirgizstan, Korea, Krasnoyarsk, Krym, Kuril Is., Lebanon-Syria, Madagascar, Magadan, Maine, Manchuria, Manitoba, Maryland, Massachusetts, Mexico Northeast, Mexico Northwest, Michigan, Minnesota, Missouri, Mongolia, Montana, Myanmar, Nebraska, Nepal, Netherlands, Nevada, New Brunswick, New Hampshire, New Jersey, New Mexico, New York, Newfoundland, North Carolina, North Caucasus, North Dakota, North European Russia, Northwest European Russia, Northwest Territories, Norway, Nova Scotia, Ohio, Oklahoma, Ontario, Oregon, Pakistan, Pennsylvania, Poland, Portugal, Primorye, Prince Edward I., Qinghai, Québec, Rhode I., Romania, Rwanda, Sakhalin, Sardegna, Saskatchewan, South Dakota, South European Russia, Spain, Sri Lanka, Sweden, Switzerland, Tadzhikistan, Taiwan, Tanzania, Tennessee, Transcaucasus, Turkey, Turkey-in-Europe, Tuva, Ukraine, Utah, Uzbekistan, Vermont, Vietnam, Virginia, Washington, West Himalaya, West Siberia, West Virginia, Wisconsin, Wyoming, Xinjiang, Yakutskiya, Yugoslavia, Yukon |
| *Poa anceps*† | Kermadec Is., New Zealand North, New Zealand South |
| *Poa leptoclada** | Burundi, Cameroon, Congo, Eritrea, Ethiopia, Gulf of Guinea Is., India, Kenya, KwaZulu-Natal, Lesotho, Malawi, Rwanda, Saudi Arabia, Somalia, Sudan, Tanzania, Uganda, Yemen, Zaïre, Zimbabwe |
| *Poa schimperiana** | Burundi, Cameroon, Congo, Ethiopia, Gulf of Guinea Is., Kenya, Malawi, Nigeria, Rwanda, Saudi Arabia, Sudan, Tanzania, Uganda, Yemen, Zaïre |
| *Pseudobromus africanus* | Burundi, Cape Provinces, Kenya, KwaZulu-Natal, Malawi, Mozambique, Northern Provinces, Rwanda, Sudan, Tanzania, Uganda, Zambia |

| | |
|---|---|
| *Secale cereale* | Turkey |
| *Setaria megaphylla** | Angola, Benin, Burundi, Cameroon, Cape Provinces, Central African Republic, Congo, Eritrea, Ethiopia, Gabon, Gambia, Ghana, Guinea, Guinea-Bissau, Gulf of Guinea Is., Ivory Coast, Kenya, KwaZulu-Natal, Liberia, Madagascar, Malawi, Mauritius, Mozambique, Nigeria, Northern Provinces, Rwanda, Réunion, Saudi Arabia, Senegal, Sierra Leone, Somalia, Sudan, Swaziland, Tanzania, Togo, Uganda, Yemen, Zambia, Zaïre, Zimbabwe |
| *Sorghum halepense* | Afghanistan, Algeria, Canary Is., Cape Verde, Chad, Cyprus, East Aegean Is., Egypt, Gulf States, India, Iran, Iraq, Kazakhstan, Kirgizstan, Kuwait, Laos, Lebanon-Syria, Libya, Madeira, Morocco, Myanmar, Nicobar Is., North Caucasus, Oman, Pakistan, Palestine, Saudi Arabia, Sri Lanka, Tadzhikistan, Thailand, Transcaucasus, Tunisia, Turkey, Turkmenistan, Uzbekistan, Vietnam, West Himalaya |
| *Sporobolus michauxianus*[†] | Alberta, Arkansas, British Columbia, Colorado, Connecticut, Delaware, District of Columbia, Idaho, Illinois, Indiana, Iowa, Kansas, Kentucky, Louisiana, Maine, Manitoba, Maryland, Massachusetts, Michigan, Minnesota, Missouri, Montana, Nebraska, New Brunswick, New Hampshire, New Jersey, New Mexico, New York, Newfoundland, North Carolina, North Dakota, Northwest Territories, Ohio, Oklahoma, Ontario, Oregon, Pennsylvania, Prince Edward I., Québec, Rhode I., Saskatchewan, South Dakota, Tennessee, Texas, Utah, Vermont, Virginia, Washington, West Virginia, Wisconsin, Wyoming |
| *Themeda triandra** | Algeria, Andaman Is., Angola, Assam, Bangladesh, Botswana, Burkina, Burundi, Cameroon, Cape Provinces, Cape Verde, Caprivi Strip, Central African Republic, Chad, China North-Central, China South-Central, China Southeast, Djibouti, East Himalaya, Egypt, Eritrea, Ethiopia, Free State, Ghana, Guinea, Hainan, India, Inner Mongolia, Ivory Coast, Japan, Jawa, Kenya, Korea, KwaZulu-Natal, Laos, Lebanon-Syria, Lesotho, Lesser Sunda Is., Madagascar, Malawi, Malaya, Mali, Maluku, Morocco, Mozambique, Myanmar, Namibia, Nansei-shoto, Nepal, New Guinea, New South Wales, Nicobar Is., Nigeria, Northern Provinces, Northern Territory, Pakistan, Philippines, Queensland, Rwanda, Saudi Arabia, Senegal, Socotra, Somalia, South Australia, Sri Lanka, Sudan, Sulawesi, Sumatera, Swaziland, |

| | |
|---|---|
| | Taiwan, Tanzania, Tasmania, Thailand, Tibet, Tunisia, Turkey, Uganda, Victoria, Vietnam, West Himalaya, Western Australia, Yemen, Zambia, Zaïre, Zimbabwe |
| *Tripidium arundinaceum* | Andaman Is., Assam, Bangladesh, Borneo, Cambodia, China North-Central, China South-Central, China Southeast, East Himalaya, Hainan, India, Jawa, Korea, Laos, Lesser Sunda Is., Malaya, Maluku, Myanmar, Nepal, New Guinea, Philippines, South China Sea, Sri Lanka, Taiwan, Thailand, Tibet, Vietnam, West Himalaya |
| *Urochloa brizantha*† | Angola, Benin, Botswana, Burkina, Burundi, Cameroon, Cape Provinces, Central African Republic, Chad, Congo, Eritrea, Ethiopia, Free State, Gabon, Gambia, Ghana, Guinea, Ivory Coast, Kenya, KwaZulu-Natal, Madagascar, Malawi, Mali, Mozambique, Namibia, Nigeria, Northern Provinces, Rwanda, Seychelles, Sierra Leone, Somalia, Sudan, Swaziland, Tanzania, Uganda, Yemen, Zambia, Zaïre, Zimbabwe |
| *Zea mays* | Guatemala, Mexico Central, Mexico Southwest |

Table S3. Metadata for the assemblages of grass pollen grains extracted from surface sediments of Mt. Kenya lakes.

| Site | Elevation (m) | Number of pollen grains |
|---|---|---|
| Lake Nkunga | 1820 | 50 |
| Rumuiku Swamp | 2154 | 50 |
| Lake Rutundu | 3078 | 50 |
| Lake Ellis | 3474 | 50 |
| Small Hall Tarn | 4500 | 50 |
| Simba Tarn | 4585 | 49 |

Table S4. Metadata for the 24 assemblages of fossil grass pollen extracted from the 25,000-year sediment record of Lake Rutundu, including the number of pollen grains analyzed from each time window. Calibrated ages (cal. yr BP) are derived from the age model of (24), based on $^{14}C$ dating of bulk organic carbon. We rounded age values to the nearest century to account for analytical and age model uncertainty.

| Core Depth (cm) | Calibrated Age (cal. yr BP) | Rounded Age (cal. yr BP) | Number of grass pollen grains |
|---|---|---|---|
| 20 | 252.922 | 300 | 49 |
| 39 | 1006.79 | 1000 | 51 |
| 71 | 2446.333 | 2400 | 54 |
| 124 | 4497.479 | 4500 | 51 |
| 134 | 5001.556 | 5000 | 51 |
| 163 | 6496.673 | 6500 | 51 |
| 173 | 7022.491 | 7000 | 50 |
| 195 | 8195.662 | 8200 | 51 |
| 236 | 9281.874 | 9300 | 49 |
| 256 | 10554.34 | 10600 | 50 |
| 263 | 11004.19 | 11000 | 49 |
| 286 | 12497.5 | 12500 | 50 |
| 298 | 13285.34 | 13300 | 50 |
| 321 | 14230.91 | 14200 | 50 |
| 335 | 14995.26 | 15000 | 50 |
| 375 | 17204.36 | 17200 | 50 |
| 397 | 18434.3 | 18400 | 49 |
| 423 | 19520.83 | 19500 | 50 |
| 432 | 20040.09 | 20000 | 48 |
| 453 | 21257.91 | 21300 | 49 |
| 470 | 22249.94 | 22200 | 51 |

| | | | |
|---|---|---|---|
| 483 | 23012.15 | 23000 | 51 |
| 516 | 24014.17 | 24000 | 50 |
| 531 | 25000.81 | 25000 | 50 |

Table S5. Five-fold cross-validation classification accuracy for the holistic CNN (H-CNN), patch CNN (P-CNN), and fused model (FM), with mean and standard deviation (s.d.).

| Split Number | Classification Accuracy (%) | | |
|---|---|---|---|
| | **H-CNN** | **P-CNN** | **FM** |
| 1 | 62.53 | 73.39 | 73.64 |
| 2 | 59.69 | 71.58 | 74.16 |
| 3 | 61.50 | 73.64 | 76.23 |
| 4 | 60.47 | 71.32 | 72.87 |
| 5 | 62.79 | 72.61 | 72.87 |
| **Mean** | 61.40 | 72.51 | 73.95 |
| **s.d.** | 1.33 | 1.04 | 1.38 |
